# Growth factor-induced desialylation for the fast control of endocytosis

**DOI:** 10.1101/2023.09.12.557183

**Authors:** Ewan MacDonald, Alison Forrester, Cesar A. Valades-Cruz, Thomas D. Madsen, Joseph H. R. Hetmanski, Estelle Dransart, Yeap Ng, Rashmi Godbole, Ananthan Akhil Shp, Ludovic Leconte, Valérie Chambon, Debarpan Ghosh, Alexis Pinet, Dhiraj Bhatia, Bérangère Lombard, Damarys Loew, Martin R. Larson, Hakon Leffler, Dirk J. Lefeber, Henrik Clausen, Patrick Caswell, Massiullah Shafaq-Zadah, Satyajit Mayor, Roberto Weigert, Christian Wunder, Ludger Johannes

## Abstract

It is commonly assumed that the glycan makeup of glycoproteins that reach the cell surface is final and static. Here, we challenge this notion by the discovery of a molecular switch that induces acute and reversible changes of glycans on the plasma membrane. We demonstrate that within minutes, the epidermal growth factor triggers the galectin-driven endocytosis of cell surface glycoproteins, such as integrins, that are key regulators of cell adhesion and migration. The onset of this process, mediated by the Na^+^/H^+^ antiporter NHE-1 and the neuraminidases Neu1/3, requires the pH-triggered enzymatic removal of sialic acids whose presence otherwise prevents galectin binding. Desialylated glycoproteins are then retrogradely transported to the Golgi apparatus where their glycan makeup is reset, and their function is repurposed to regulate EGF-dependent invasive cell migration. Glycosylation at the cell surface thereby emerges as a dynamic and reversible regulatory post-translational modification that controls a highly adaptable trafficking pathway.

## Introduction

Growth factors engage cell surface receptors and trigger signaling events with wide cellular ramifications (Avraham and Yarden, 2011; Lemmon and Schlessinger, 2010), including effects on the glycosylation machinery and the glycosylation state of *de novo* synthesized proteins (Gao et al., 2021; Gilmour et al., 2013; Marsico et al., 2018; Palmisano et al., 2011; Stanley, 2011; Takahashi et al., 2016; Zhao et al., 2008). The possibility that growth factors affect glycans at the plasma membrane has been largely overlooked. The glycan makeup of glycoproteins that reach the cell surface is indeed generally considered to be set. Nonetheless, we here present a novel regulatory circuit based on growth factor-induced acute and reversible changes in glycans of cell surface glycoproteins which in turn repurposes their functions.

We describe the discovery of a signal transduction pathway that links growth factor signaling to the modulation of galectin-driven clathrin-independent endocytosis through the triggered removal of terminal sialic acid (Sia) residues from N-glycans on cargo client proteins. In addition to the well explored clathrin pathway (Kaksonen and Roux, 2018; Kirchhausen et al., 2014; McMahon and Boucrot, 2011; Sorkin, 2004), mechanisms for building endocytic pits without clathrin have emerged more recently (Doherty and McMahon, 2009; Johannes et al., 2015; Renard and Boucrot, 2021; Sandvig et al., 2018; Sigismund et al., 2021). One of the drivers are galectins, a large family of proteins (Cummings et al., 2022; Johannes et al., 2018) whose glycan-binding properties are greatly influenced by the degree of sialylation and the type of Sia linkage (α2-3 or α2-6) (Leffler and Barondes, 1986; Stowell et al., 2008; Zhuo and Bellis, 2011). Therefore, Sia capping of glycoproteins and glycolipids enhances or inhibits galectin binding interactions with wide biological effects (Läubli and Varki, 2020; Varki and Gagneux, 2012). Amongst others, galectins tune the cell surface dynamics of glycoproteins through lattice formation (Nabi et al., 2015; Partridge et al., 2004; Rabinovich et al., 2007) and glycolipid-lectin (GL-Lect) driven endocytosis (Johannes et al., 2016; Lakshminarayan et al., 2014; Renard et al., 2020) in an intertwined manner that requires further in-depth investigation (Mathew and Donaldson, 2018).

We show that signal transduction between the archetypical epidermal growth factor receptor (EGFR) and the membrane sialidases (neuraminidases, Neu) that catalyze desialylation, involves a previously unrecognized acidification-dependent reaction at the cell surface. Among others, we identify integrins as substrates of EGF-induced desiaylation. Integrins are heterodimeric transmembrane proteins composed of one α and one β subunit that bind directly to the extracellular matrix (Bridgewater et al., 2012; Marsico et al., 2018; Moreno-Layseca et al., 2019). ST6GAL1-catalyzed sialic acid modification (i.e., α2-6 sialylation) has been associated in physiological and pathological conditions with integrin activation, fibronectin affinity modulation, and cell migration (Hou et al., 2016; Pan and Song, 2010; Pretzlaff et al., 2000; Seales et al., 2005). Of note, endocytic trafficking of integrins has been shown to be important for their activities (Moreno-Layseca et al., 2019).

Here, we demonstrate that following EGF-induced desialylation, integrins are transported via the retrograde route to the Golgi apparatus, whereupon their glycan makeup is reset by resialylation and their functions repurposed by polarized secretion to the leading edge to support invasive cell migration. Based on these findings, we propose a model, termed the desialylation glycoswitch, by which extracellular cues are relayed to galectin-driven endocytic processes and transduced into pathophysiological responses through the acute remodeling of cell surface glycans.

## Results

### Growth factors trigger desialylation

Galectin-3 (Gal3) is the best characterized driver of an uptake process that has been termed glycolipid-lectin (GL-Lect) driven endocytosis (Johannes et al., 2016). For this, oligomerization competent Gal3 reorganizes glycoproteins and glycosphingolipids in a way that triggers the biogenesis of tubular endocytic pits from which clathrin-independent endocytic carriers (CLICs) emerge (Johannes et al., 2016; Renard and Boucrot, 2021).

To study if the GL-Lect process is regulated by growth factor signaling, we measured exogenous Gal3 binding to MDA-MB-231 breast cancer cells incubated in the presence of saturating concentrations (100 ng/mL) of a panel of growth factors (Figure 1A). These incubations were performed at 4 °C where endocytosis is inhibited but growth factors continue to signal (de Melker et al., 2001; Stang et al., 2004). Gal3 binding to cells was significantly increased after treatment with several growth factors, including EGF and TGFα, both ligands of EGFR (Figures 1A and 1B).

**Figure 1.**
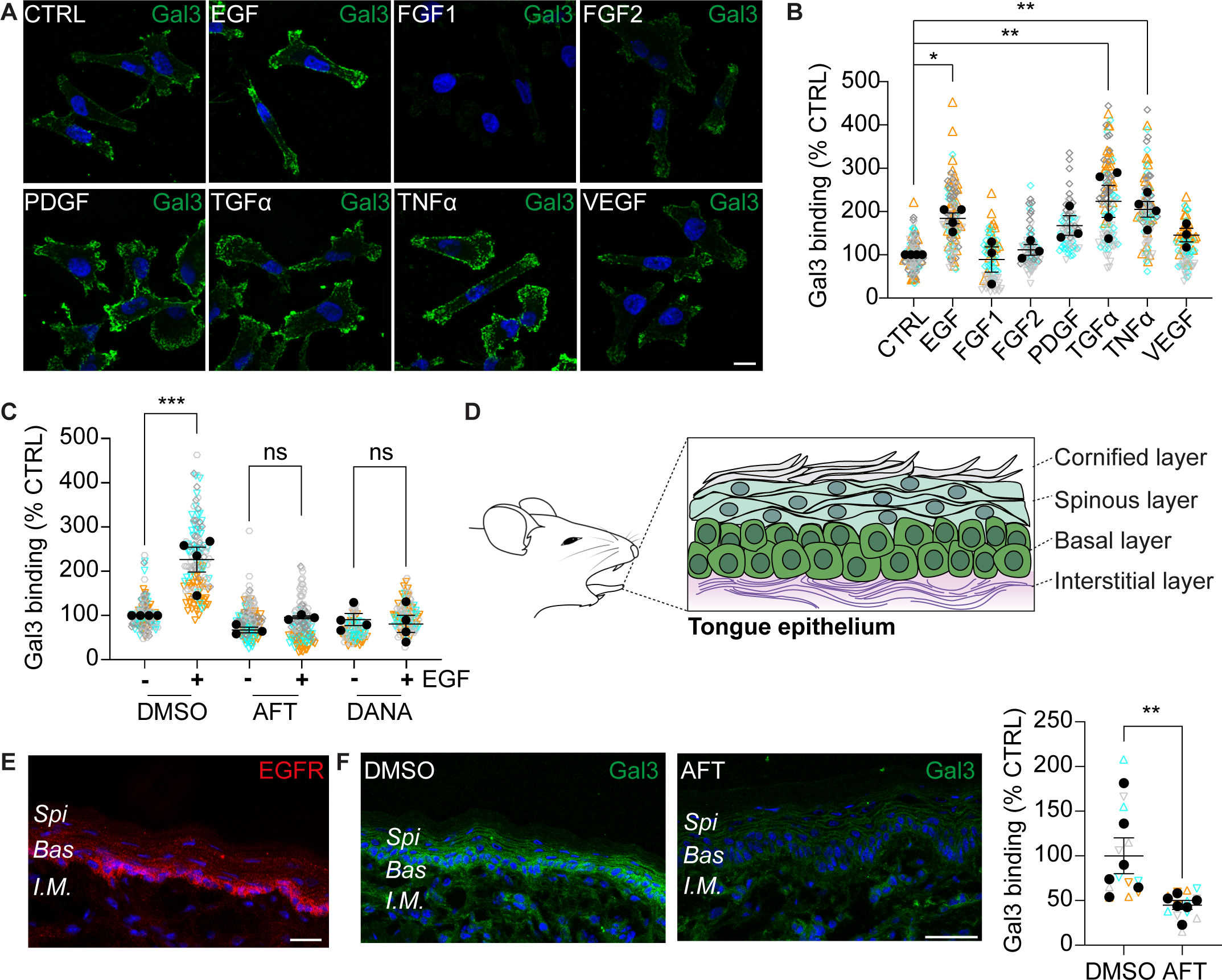
EGF modulates Gal3 binding to cells. (A) Growth factor screening. Serum starved MDA-MB-231 cells were incubated for 1 h on ice with Alexa488-labeled Gal3 in the presence or absence of 100 ng/mL of the indicated growth factors. (B) Fluorescence intensity was determined for 4 independent experiments as in (A) (≈ 100 cells per condition). One-way ANOVA with Dunnett’s multiple comparison test. (C) Experiment as in (A) with EGF under conditions of afatinib (AFT) or DANA treatment. 4 independent experiments (≈ 150 cells per condition). One-way ANOVA with Dunnett’s multiple comparison test. (D) Simplified scheme of the epithelial architecture of the mouse tongue. (E) EGFR (red) immunolabeling on tissue section of the ventral part of the mouse tongue. (F) Gal3 localization on sections of tongues from mice that had been treated or not with AFT. The mean intensity of Gal3 labeling was analyzed at the level of the basal layer on 6 mice per condition. Mann-Whitney test. Scale bars = 10 µm (A) or 45 µm (E,F). In (A,E,F): Nuclei in blue. In (B,C,F): Means ± SEM. Means from separate experiments are indicated by solid dots, and measurements of individual cells have different colored symbols for each experiment. ns = p > 0.05, * p ≤ 0.05, ** p < 0.01, *** p < 0.001.

The stimulation of Gal3 binding was observed with as little as 1 ng/mL of EGF in MDA-MB-231 cells, illustrating the exquisite sensitivity of the system (Figure S1A). This effect was also found in HN12 tongue squamous cell carcinoma cells and mouse embryonic fibroblasts (MEFs) (Figures S1B and S1C). However, in NR6 mouse fibroblasts that lack EGFR (Pruss and Herschman, 1977), the EGF-stimulated Gal3 binding was only observed upon ectopic EGFR expression (Figure S1D). Furthermore, the pretreatment of MDA-MB-231 (Figure 1C) or HN12 (Figure S1B) cells with afatinib (AFT), an irreversible EGFR tyrosine kinase inhibitor (Yu and Pao, 2013), prevented the EGF-induced Gal3 binding.

Next, we determined whether the EGFR-dependent stimulation of Gal3 binding occurred under physiological conditions. Based on the results in the tongue squamous cell carcinoma-derived HN12 cells (Figure S1B), we chose the oral cavity and in particular the tongue epidermis (Figure 1D), since it is exposed to high levels of EGF (1 µg/ml) secreted into the saliva in mice (Byyny et al., 1974), and it shows high expression of EGFR in the basal layer (Figure 1E). Notably, we found that AFT administration to live animals, and not the vehicle, inhibited EGFR autophosphorylation on tyrosine 1068 (Figure S1E) and significantly reduced the binding of Gal3 to the basal layer (Figure 1F).

Altogether, these findings demonstrate that EGFR and its tyrosine kinase activity are required for EGF-induced Gal3 binding to cells *in vitro* and in the complex environment of a living organism.

Gal3 binding is partly permissive to α2-3-linked Sia capping of glycans but blocked by α2-6-linked Sia (Leffler and Barondes, 1986; Stowell et al., 2008; Zhuo and Bellis, 2011) (Figure 2A). We therefore tested whether the EGF-induced increase of Gal3 binding to cells was dependent on the enzymatic removal of Sia (desialylation). For this, we used the compound, 2-deoxy-2-3-didehydro-N-acetylneuraminic acid (DANA), which is a known inhibitor of endogenous neuraminidases (Neu1-4) that desialylate glycans (Glanz et al., 2018). Pretreatment of MDA-MB-231 cells (Figure 1C), HN12 cells (Figure S1B) or MEFs (Figure S1C) with DANA resulted in failure of EGF to induce Gal3 binding. This was not due to an impairment in EGFR signaling under these conditions, as EGF-induced phosphorylation of extracellular signal-regulated kinases 1/2 (ERK1/2) and ribosomal S6 kinases (RSK) was not affected by incubation of cells with DANA (Figures S1G and S1H). Treatment with sialostatin (STI, also called SiaFNEtoc), a general inhibitor of cellular sialylation (Moons et al., 2022) (Figure 2A), augmented basal Gal3 binding and abolished the EGF-induced increase (Figures S1I and S1J).

**Figure 2.**
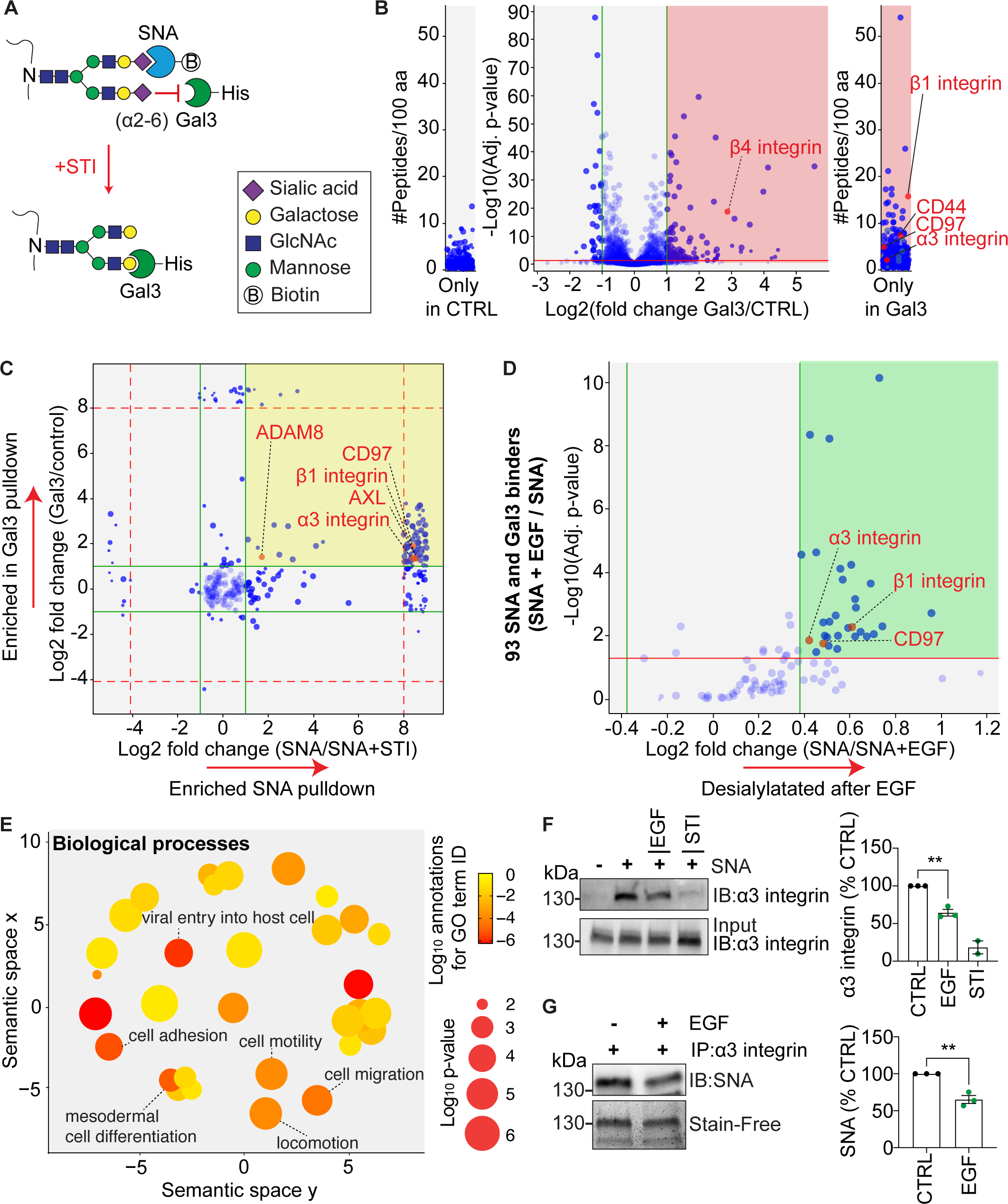
EGF induces desialylation. (A) Schematics of interaction between ⍰2-6 Sia-modified glycans and Gal3 or SNA. (B) Identification of Gal3 interactors. MDA-MB-231 cells were incubated for 1 h on ice with Gal3-His. Pulldowns were subjected to quantitative mass spectrometry analysis. Red quadrant shows significantly enriched Gal3 binders. (C) Identification of Gal3 binders that are also SNA interactors. MDA-MB-231 cells, either untreated or treated with STI, were incubated for 1 h on ice with SNA-biotin. Pulldowns were analyzed by quantitative mass spectrometry. 574 proteins from 5 independent experiments were common to Gal3 (from B) and SNA (from C) data sets. 93 of these were enriched with high confidence for both (yellow quadrant). (D) Volcano plot of these 93 high confidence interactors as to their propensity to undergo EGF-induced desialylation (experiment as in (C) in the presence of 100 ng/mL of EGF). 30 responders are highlighted in the green quadrant. Note the presence of ⍰3β1 integrin and CD97 amongst the hits. (E) Cluster analysis of the summarized list of “biological processes” gene ontology terms associated with the 30 EGF-sensitive Gal3 and SNA interactors. (F) Validation of proteomics experiments by SNA pulldown. SNA-biotin pulldown was performed as in (B) in the presence or absence of STI or EGF. Samples were analyzed by Western blotting against α3 integrin. Quantifications from 3 independent experiments. One-way ANOVA with Tukey’s multiple comparison test. (G) Validation of proteomics experiments by α3 integrin immunoprecipitation. MDA-MB-231 cells were incubated for 1 h on ice anti-α3 integrin antibodies in the presence or absence of EGF. After pulldown, samples were analyzed by Western blotting using SNA-biotin. Quantifications from 3 independent experiments. Two tailed unpaired t-test. In (F,G): ** p < 0.01, *** p < 0.001.

All these findings are consistent with the notion that the effect of EGF on Gal3 binding to plasma membrane proteins is directly dependent on the presence of Sia residues and their removal.

To identify the cell surface glycoproteins that undergo EGF-induced desialylation, we employed a multi-step lectin-enriched mass spectrometry-based approach. First, we demonstrated that binding of the α2-6 Sia linkage-specific *Sambucus nigra* (SNA) lectin (Figure 2A) to MDA-MB-231 cells was reduced after EGF treatment (Figure S1K). Interestingly, EGF treatment did not measurably affect binding of the α2-3 linkage-specific lectin-like Lectenz reagent (Figure S1L), suggesting that predominantly α2-6 Sia capped glycans are being desialylated upon EGF stimulation to serve for efficient Gal3 binding. This is in agreement with the published notion that only α2-6-linked Sia blocks glycan recognition by Gal3 (Leffler and Barondes, 1986; Stowell et al., 2008; Zhuo and Bellis, 2011) (Figure 2A).

Second, we identified Gal3 interactors at the surface on MDA-MB-231 cells (green quadrant in Figure 2B). Third, we used cell surface-bound SNA to identify α2-6 sialylated plasma membrane proteins (STI as a control for total sialylation), and then cross referenced these with the list of Gal3 interactors (Figure 2C, Table S1). We thereby identified 93 proteins that were high confidence interactors of both, Gal3 and SNA (yellow quadrant in Figure 2C, Table S1). 30 of these 93 high confidence interactors were significantly desialylated upon EGF treatment (green quadrant in Figure 2D, Table S2). Gene ontology analysis on these 30 hits revealed cell adhesion, migration, and locomotion as some of the most highly enriched terms (Figure 2E).

To further validate the proteomics data we focused amongst the 30 hits on the cell adhesion and migration factor α3β1 integrin, as integrins undergo extensive endocytic trafficking (Jones et al., 2006; Moreno-Layseca et al., 2019) and have been identified as substrates for both, Gal3-driven internalization (Furtak et al., 2001; Lakshminarayan et al., 2014) and retrograde transport in relation to persistent cell migration (Shafaq-Zadah et al., 2016). Pulldown and Western blotting experiments confirmed that the α3 chain was indeed desialylated upon EGF treatment of MDA-MB-231 cells (Figures 2F and 2G).

Therefore, we established that EGF acutely triggers desialylation of select plasma membrane proteins including those involved in cell adhesion and migration.

### EGF-induced desialylation requires Neu1 and Neu3 and is abrogated in NEU1 deficient patients

To elucidate the mechanism underlying the EGFR-induced desialylation, we first employed a siRNA-based screen for the four endogenous desialylation enzymes Neu1-4 (Monti et al., 2010). Knockdown of Neu1 and Neu3, previously reported to be localized to the plasma membrane (Maurice et al., 2016; Monti et al., 2000; Monti et al., 1999; Seyrantepe et al., 2004; Zanchetti et al., 2007), led to a loss of EGF-induced Gal3 binding to the cell surface (Figure S2A). We confirmed these results using a CRISPR-based knockout approach in MDA-MB-231 cells (Figures 3A, S2B and S2C). Importantly, autophosphorylation of EGFR was not affected in *NEU1* and *NEU3* knockout cell lines (Figure S2D), suggesting that neuraminidase activity is required for Gal3 binding downstream of EGFR signaling.

**Figure 3.**
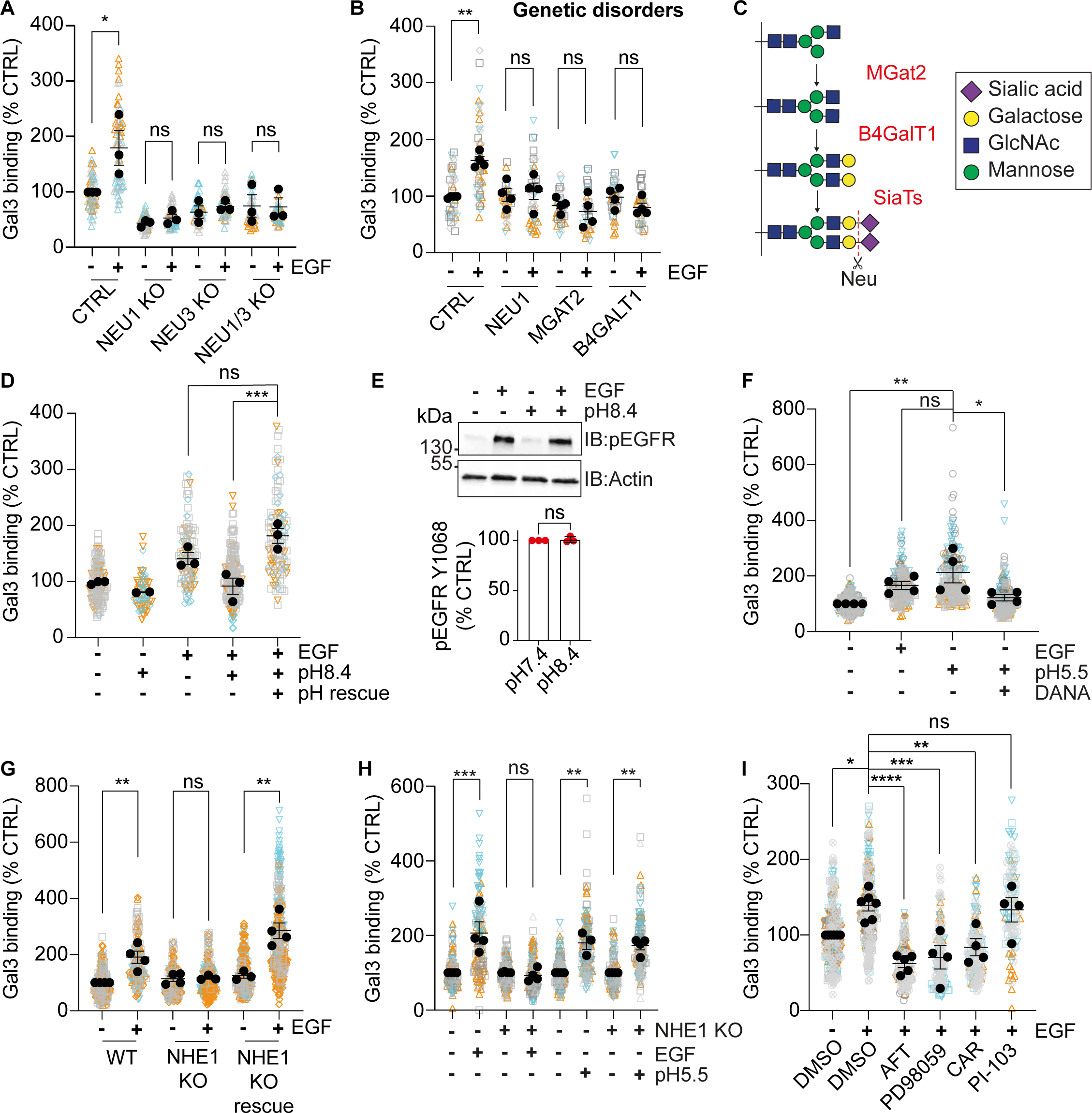
NHE1, Neu1/3, and acidification are required for EGF-induced desialylation. (A) Neuraminidases. Serum starved wildtype MDA-MB-231 cells or CRISPR-based knockouts of *NEU1*, *NEU3* or *NEU1/3* were incubated for 1 h on ice with Alexa-488 coupled Gal3 in the presence or absence of EGF. Fluorescence signals were quantified from 3 independent experiments (≈ 75 cells per condition). One-way ANOVA with Tukey’s multiple comparison test. (B) Experiment as in (A) with patient skin fibroblasts harboring inactivating mutations in the indicated enzymes. 4 independent experiments (≈ 100 cells per condition). One-way ANOVA with Tukey’s multiple comparison test. (C) Schematics of N-glycosylation pathway. (D) pH quenching effect. Serum starved MDA-MB-231 cells were incubated for 1 h on ice at pH 7.4 or pH 8.4 in the presence or absence of EGF, fixed, and incubated at neutral pH with Alexa-488 coupled Gal3. Quantifications as in (A) on 3 independent experiments (≈ 100 cells per condition). One-way ANOVA with Dunnett’s multiple comparison test. (E) Cells were treated as in (D), lysed after EGF stimulation, and analyzed by Western blotting against EGFR Y1068. 3 independent experiments. Unpaired t-test. (F) pH triggering effect. Serum starved MDA-MB-231 cells were incubated for 1 h on ice in pH 5.5 or pH 7.4 buffers with or without EGF and/or DANA, and then for 1 h on ice at neutral pH with Alexa-488 coupled Gal3. Quantifications as in (A) on 4 independent experiments (≈ 100 cells per condition). One-way ANOVA with Dunnett’s multiple comparison test. (G) Role of NHE1, CRISPR. Serum starved wild-type MDA-MB-468 cells, CRISPR-based knockouts of NHE1, and NHE1-GFP rescue cells were incubated and quantified as in (A) on 4 independent experiments (≈ 100 cells per condition). One-way ANOVA with Dunnett’s multiple comparison test. (H) pH 5.5 buffer can rescue NHE knockout cells. Serum starved MDA-MB-231 wildtype and NHE1 knockout cells were incubated for 1 h on ice in pH 5.5 or pH 7.4 buffers with or without EGF. Quantifications as in (A) on 4 independent experiments and normalized to controls (≈ 100 cells per condition). One-way ANOVA with Dunnett’s multiple comparison test. (I) Role of NHE1, inhibitors. MDA-MB-231 cells were pretreated for 30 min at 37 °C with the indicated inhibitors or DMSO (CTRL) in serum free media before incubation in the continued presence of the inhibitors for 1 h on ice with Alexa-488 coupled Gal3 in the presence or absence of EGF. Quantifications as in (A) on 5 independent experiments (≈150 cells per condition). One-way ANOVA with Dunnett’s multiple comparison test. Means ± SEM are shown in this figure. Except for (E): Means from separate experiments are indicated by solid dots, and measurements of individual cells have different colored symbols for each experiment. ns = p > 0.05, * p ≤ 0.05, ** p < 0.01, *** p < 0.001, **** p < 0.0001.

Deficiency in *NEU1* is found in the rare lysosomal storage disease type 1 sialidosis (Bonten et al., 1996; Sphranger et al., 1977). We found here that *NEU1* deficiency abolished EGF-induced Gal3 binding to patient-derived fibroblasts, while fibroblasts from healthy donors showed the expected increase of Gal3 binding (Figure 3B). We further tested fibroblasts from donors with genetic deficiencies in the upstream biosynthetic pathway leading to Sia capped N-glycans, including the MGAT2 and B4GALT1 enzymes (Figure 3C). These cells also did not show enhanced Gal3 binding after EGF stimulation (Figure 3B). While the glycosylation of β1 integrin was altered in *MGAT2* or *B4GALT1*-deficient cells (Figure S2E), the EGFR-dependent MAPK signaling remained intact in all the tested deficiency conditions (Figure S2F), further supporting that loss of EGF-induced Gal3 binding on mutant cells was not due to a general perturbation of EGFR signaling.

These findings provide strong genetic evidence for the link between EGFR signaling and Neu1/3-mediated desialylation of plasma membrane proteins leading to increased Gal3 binding.

### Acidification-based mechanism for desialylation at the plasma membrane

We next set out to identify the signaling pathway linking EGFR and Neu1/3. While it has been reported that Neu1 can be phosphorylated downstream of growth factor receptors in lymphocytes (Lukong et al., 2001), we failed to detect EGF-induced changes in the phosphorylation status of Neu1 or Neu3 in MDA-MB-231 cells. These enzymes, which exhibit a limited activity at the pH of the extracellular medium (pH 7.4), work optimally at around pH 5.5 (Albohy et al., 2010). Therefore, we hypothesized the existence of a mechanism of transient acidification at the plasma membrane. As the hydroxide is the fastest agent to buffer protons, we tested if step wise incubation of MDA-MB-231 cells in an alkaline buffer (incubation with EGF at pH 8.4, followed by incubation with Gal3 at neutral pH) affected the EGF-induced Gal3 binding, and remarkably found that high pH abrogated the EGF effect (Figure 3D). In this condition, EGFR activation still occurred, as shown by the fact that EGF-induced autophosphorylation of EGFR on tyrosine 1068 was not affected (Figure 3E), as previously reported (Nunez et al., 1993). Furthermore, the abrogation of Gal3 binding at alkaline pH was fully reversible by restoring physiological pH (Figure 3D, pH rescue). Finally, incubating cells in pH 5.5 buffer without EGF induced a similar increase in Gal3 binding as with incubation with EGF (Figure 3F). Of note, this pH 5.5-triggered effect on Gal3 binding was strictly dependent on sialidase activity as it was abolished in the presence of DANA (Figure 3F).

These findings demonstrate that extracellular acidification is necessary and sufficient for triggering Neu1/3-mediated Gal3 binding to cells.

A clue as to how the extracellular pH might be controlled came from studies in which it was shown that EGFR induces the acidification of the extracellular milieu through the activation of the NHE1 channel, a Na^+^/H^+^ antiporter whose activity is correlated with tumor malignancy (Fliegel, 2021; Koivusalo et al., 2010; Sardet et al., 1990; Stock and Pedersen, 2017). We found that NHE1 was required for EGF-induced Gal3 binding, using MDA-MB-468 breast cancer cells with *NHE1* knockout and re-expression rescue (Figure 3G). Notably, pH 5.5 incubation stimulated Gal3 binding even on *NHE1* knockout cells (Figure 3H), thus positioning the acidification of the extracellular medium downstream of NHE1 (Figure 3H) and upstream of Neu1/3 (DANA in Figure 3F). EGFR was also reported to stimulate NHE1 activity by phosphorylation at position S703, through signaling mediated by RAF-MEK-ERK and p90RSK kinases, and at positions S766, S770, and S771, by ERK1/2 (Takahashi et al., 1999). We found that a triple mutation in the NHE1 gene (S766A, S770A, S771A; termed SSSA) rescued the EGF-induced Gal3 binding in MDA-MB-231 cells lacking NHE1, whereas a mutation at position S703 (S703A) did not (Figure S2G). This identifies phosphorylation of NHE1 at S703 as a critical event for EGFR signaling leading to increased Gal3 binding.

Pharmacological inhibition of NHE1 using cariporide (CAR) (Scholz et al., 1993) also led to the loss of the EGF effect on Gal3 binding (Figures 3I and S2H). The involvement of the MAP kinase pathway was confirmed by pharmacological interference with MEK activity using the inhibitor PD98059 (Figures 3I and S2H). In contrast, the phosphoinositide 3-kinase inhibitor PI-103 had no effect on EGF-induced Gal3 binding (Figure 3I), indicating that the PI3K/Akt pathway is not involved.

NHE1 thereby is a key mediator between EGFR signaling and extracellular acidification leading to Neu1/3-catalyzed desialylation of cargo proteins and subsequent Gal3 binding.

Collectively, these results identify a pathway that is triggered by the activation of EGFR signaling and involves the sequential downstream activation of 1) the sodium–proton antiporter NHE1, which induces a transient acidification, 2) the neuraminidases Neu1 and Neu3 which catalyze desialylation of plasma membrane glycoproteins, and 3) Gal3 binding to these glycoproteins.

### EGF-induced desialylation triggers endocytosis

Since Gal3 is a key driver of clathrin-independent endocytic uptake according to the GL-Lect mechanism, we tested whether EGF-induced desialylation modulated α3β1 integrin endocytosis in a Gal3-dependent manner. When MDA-MB-231 cells were treated with EGF, the uptake of a fluorescently labeled anti-α3 integrin antibody was significantly increased, whereas the internalization of transferrin, a classic clathrin pathway cargo, was not affected (Figures 4A and 4B). The EGF-induced stimulation of α3 integrin uptake was lost when incubations were made in the presence of the membrane impermeable Gal3-specific inhibitor 3 (termed I3 in the current study; GB0149-03 in (Salameh et al., 2010; Stegmayr et al., 2019)) (Figure 4B), which established that the extracellular fraction of Gal3 was required for this effect. EGF also induced the uptake of β1 integrin (Figures 4C, S2I and S2J) and of the G protein coupled receptor CD97 (Figure 4D and 4E), which was also present in the list of cargoes that are desialylated upon incubation with EGF (Figure 2D). For both cargoes, this EGF-induced uptake occurred in a neuraminidase-dependent manner (Figures 4C and 4E). Furthermore, interference with EGFR tyrosine kinase activity, NHE1, and RSK kinase (BI-D1870) resulted in a loss of EGF-stimulated endocytosis of β1 integrin (Figure 4F). This highlights that the signal transmission chain between EGFR and cargo protein desialylation is involved in the stimulatory effect of EGF on Gal3-dependent α3β1 integrin and CD97 uptake. Of note, TNFα, which is amongst the growth factors that trigger Gal3 binding to cells (Figure 1B), also stimulated β1 integrin uptake in a sialidase-sensitive manner (Figure 4C), which suggests that the described desialylation mechanism may apply in signaling contexts other than EGF.

**Figure 4.**
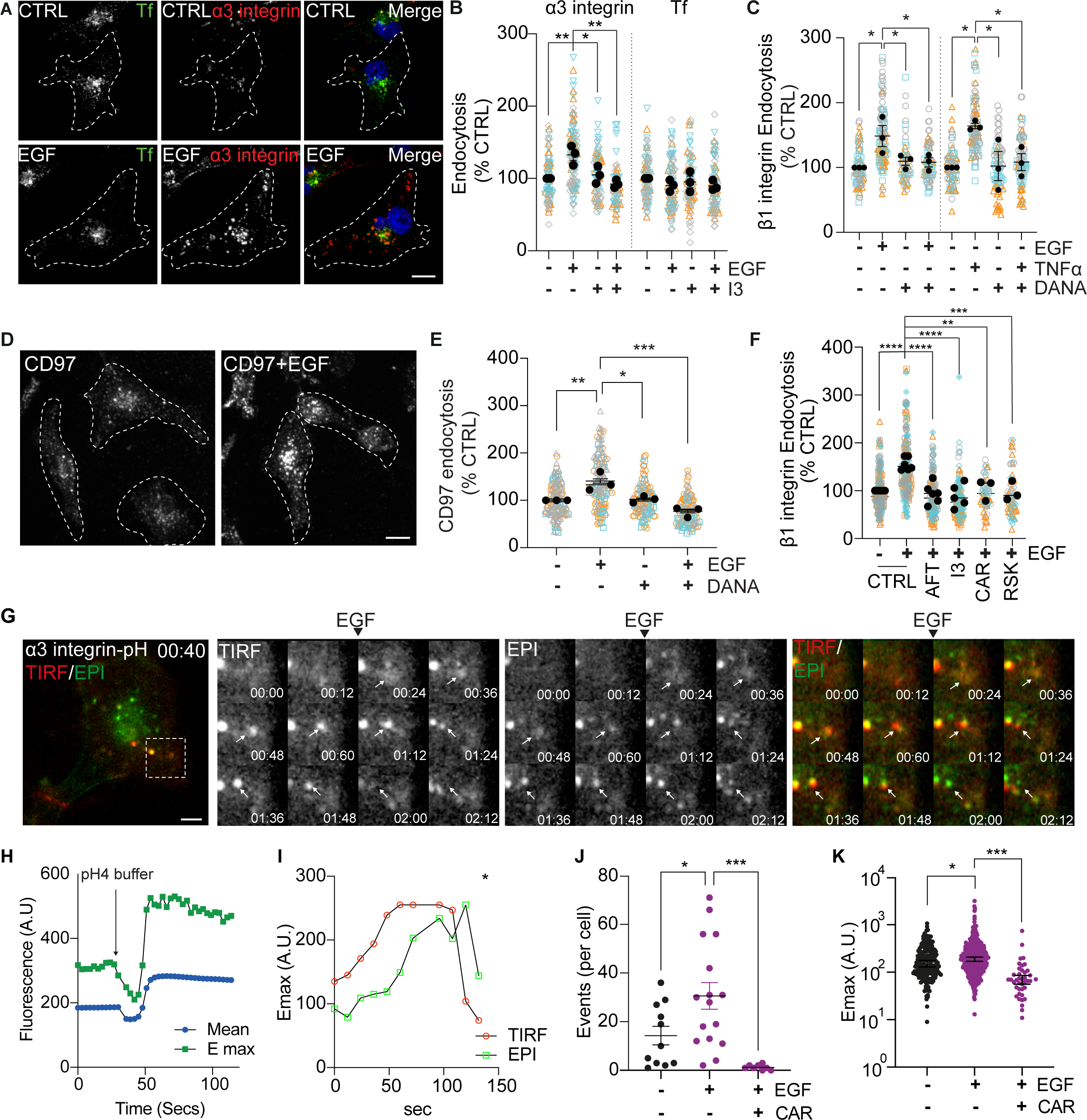
EGF-induced desialylation triggers GL-Lect driven endocytosis. (A) ⍰3 integrin uptake. Serum starved MDA-MB-231 cells were continuously incubated for 10 min at 37 °C in the presence or absence of EGF with Atto647N-labeled anti-⍰3 integrin antibody and Alexa488-transferrin (Tf). Note that ⍰3 integrin uptake was stimulated in the presence of EGF. Scale bar = 10 µm. (B) Quantification of fluorescence from 3 independent experiments (≈ 100 cells per condition) as in (A) in which cells were incubated in the presence or absence of EGF and/or the Gal3 inhibitor I3. One-way ANOVA with Dunnett’s multiple comparison test. (C-F) β1 integrin and CD97 uptake. Experiments as in (A) in which anti-β1 integrin or anti-CD47 antibodies were used in the presence or absence of the indicated inhibitors and growth factors. 3-6 independent experiments (≈ 200 cells per condition). One-way ANOVA with Dunnett’s multiple comparison test. In (D): Scale bar = 10 µm. (G) pH imaging. MDA-MB-231 cells were incubated continuously for 4 min at 37 °C with AcidiFluor-coupled anti-α3 integrin antibody, and then imaged for 10 min sequentially in TIRF and epifluorescence modalities. EGF was added after 18 sec. Note that increased fluorescence in the TIRF field (red) indicates acidification. Arrows indicate 1 representative event. Scale bar = 10 µm. (H) Dynamic pH monitoring. Acquisition of images started immediately after incubation with antibody in sequential TIRF and epifluorescence modes at 37 °C. pH 4 buffer was added at the indicated time point to document how the setup reacted. Total fluorescence (mean) and Emax are shown for a representative cell. (I) Intensity traces for the event shown by arrows in (G). * indicates where vesicle leaves frame. TIRF in red, widefield in green. (J,K) Quantification of numbers per cell (I) or of maximum intensity values (Emax) (J) for tracks that undergo a pH change as shown in (G). 3 independent experiments (≈ 15 cells counted per condition). Holm-Šídák’s multiple comparisons test. Means ± SEM are shown in this figure. In (B,C,E,F), means from separate experiments are indicated by solid dots, and measurements of individual cells have different colored symbols for each experiment. * p ≤ 0.05, ** p < 0.01, *** p < 0.001.

To directly test whether an EGF-induced transient decrease of the extracellular pH occurred in the immediate vicinity of endocytic events, anti-α3 integrin antibodies were coupled to the pH-sensitive dye AcidiFluor Orange, which is non-fluorescent at neutral pH and becomes fluorescent upon acidification. Using a microscopy setup that allows switching within seconds between total internal reflection fluorescence (TIRF) and widefield acquisitions (Figures 4G and 4H), we recorded the pH surrounding α3 integrin from locations at or near the plasma membrane to deeply internalized endocytic carriers. We found that low pH signal was first observed in the plasma membrane-proximal zone by TIRF microscopy (Figure 4I, red), upon which internalization into the full volume of cells was tracked by widefield microscopy (green). The incubation of cells with EGF increased the number (Figure 4J) and peak intensities (Figure 4K) of these events. Analysis of endocytic events by lattice light sheet microscopy (LLSM) (Figures S2K and S2L) was consistent with the conclusions that the increased fluorescence signal of AcidiFluor Orange-tagged anti-α3 integrin antibody resulted from EGF-induced acidification in the immediate environment of the cargo to which it was bound, and not from increased amounts of cargo. This EGF effect was lost in the presence of cariporide, which again strongly argued for the involvement of NHE1 in the observed acidification process (Figures 4J and 4K).

Ultrastructural studies revealed very few small (80-120 nm) or large (>120 nm) vesicular carriers containing α3 or β1 integrin (Figures 5A, S3A and S3B). In contrast, both proteins were abundantly found in short tubular and crescent shaped structures, a morphological hallmark of clathrin-independent endocytic carriers (CLICs) (Howes et al., 2010; Lakshminarayan et al., 2014). Quantification revealed an increase of β1 integrin-positive CLICs upon incubation of cells with EGF (Figure 5B).

**Figure 5.**
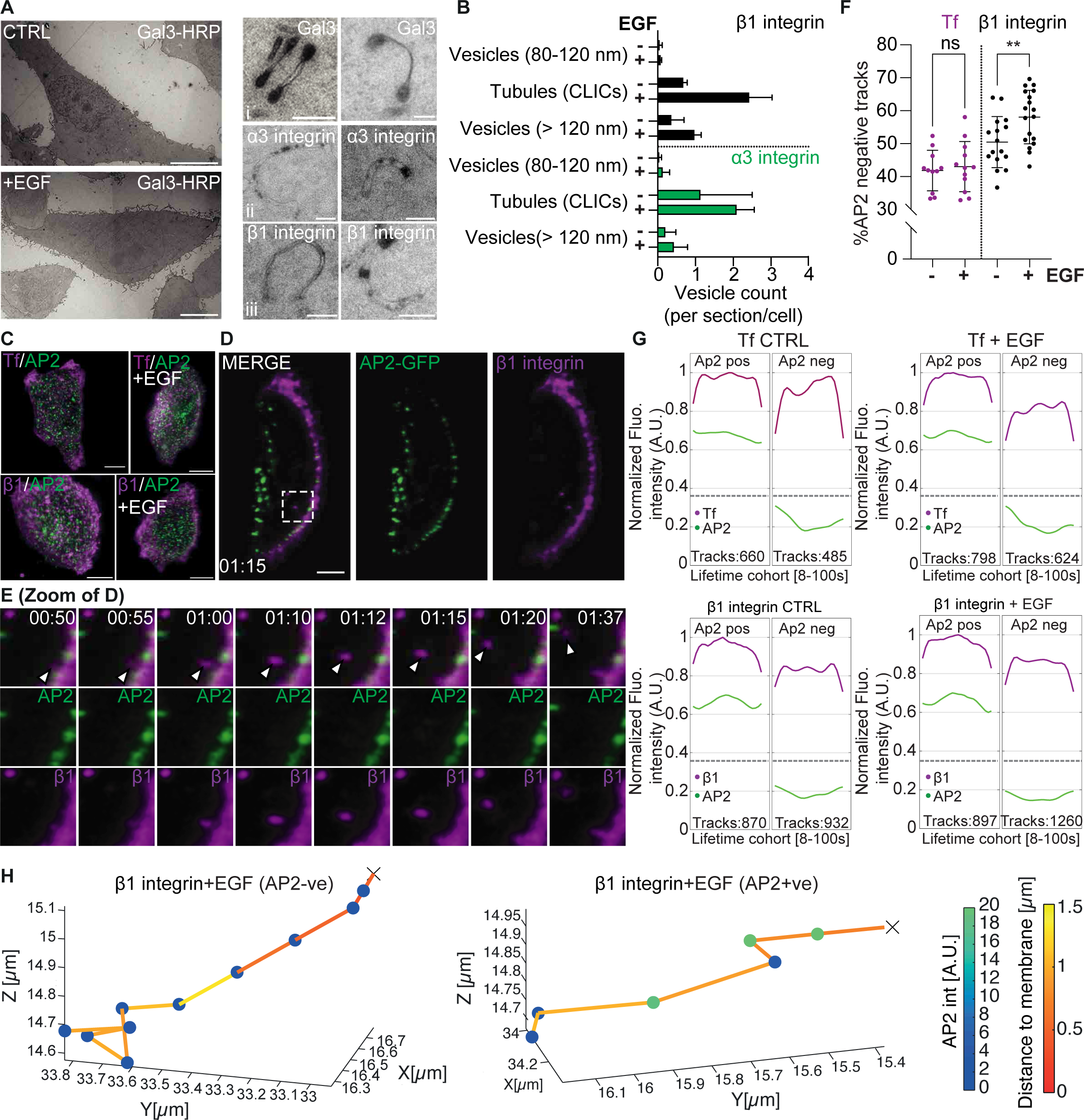
EGF-induced desialylation induces formation of clathrin-independent carriers. (A) Electron microcopy. Left: Representative overview electron micrographs of MDA-MB-231 cells that have been incubated with HRP-labeled Gal3 for 6 min at 37 °C in the presence or absence of EGF. Right: Zoomed views of CLICs from EGF-treated MDA-MB-231 cells for which the following HRP-labeled ligands were present during incubations: (i) Gal3, (ii) anti-α3 integrin antibodies, or (iii) anti-β1 integrin antibodies. Scale bars = 10 µm (images on left) and 200 nm (images on right). (B) Quantification of electron micrographs as in (A). HRP-positive structures from 2 independent experiments were counted for the indicated conditions (≈ 10 cells per condition) in a single slice per cell and categorized according to morphology. (C) Lattice light sheet microscopy. Representative maximum intensity projections of AP2-eGFP (green) expressing SUM159 cells in indicated conditions of incubation with anti-β1 integrin-Cy3 antibodies or Tf-Cy3 (magenta) in the presence or absence of EGF. Scale bars = 10 µm. (D) Central section of a SUM159 cell at 1 min 15 s into acquisition, showing an AP2-negative β1 integrin uptake event (in dotted box). Scale bar = 5 µm. (E) High magnifications panels of single frames from boxed area in (D). The AP2-negative uptake event is tracked by white arrowheads. (F) Percentage of β1 integrin and transferrin (Tf) uptake tracks from 3 independent experiments that were AP2-negative as in (C-E). For β1 integrin: 1802 tracks from 16 control cells, and 2157 tracks from 18 EGF-treated cells. For Tf: 1145 tracks from 11 control cells, and 1422 tracks from 12 EGF-treated cells. Note that the percentage of AP2-negative tracks increased upon EGF treatment only for β1 integrin uptake. Means ± SEM, unpaired t-test. (G) Average normalized intensity traces and classifications for β1 integrin and Tf uptake events from experiments in (F). Lifetime cohorts of endocytic trajectories show “dome”-shaped intensity profiles for cargoes, and co-tracking or not with AP2. Dotted lines represent background levels for fluorescence signals. (H) Selected β1 integrin tracks in EGF-treated cells. Axes show X,Y,Z positions of uptake carriers in cells, spot colors depicts AP2 intensity, line color distance to plasma membrane, and X indicates track starting points. Means ± SEM are shown in this Figure. ns = p > 0.05, ** p < 0.01.

To dynamically map endocytic uptake events for β1 integrin, we employed LLSM on gene edited triple negative SUM159 breast cancer cells expressing AP2-eGFP (Aguet et al., 2016), a key adaptor of the clathrin pathway (Kural et al., 2015). LLSM allowed for 3D imaging of live cells at high acquisition frequency (Figures 5C, 5D and 5E and Movies S1-4). Using advanced image-processing techniques (Renard et al., 2020), we detected and tracked diffraction-limited β1 integrin spots, and extracted data from single endocytic events per cell, including distance from the plasma membrane and fluorescence intensity of β1 integrin and AP2-eGFP (Figures 5F, 5G and 5H). To define whether β1 integrin was endocytosed together with clathrin, we measured whether tracks were positive or negative for AP2 (Figures 5D and 5E, Movie S5). While a minority of transferrin endocytic tracks were AP2-negative, β1 integrin endocytic tracks were equally distributed between AP2-negative and positive categories (Figures 5F and 5G). Only for β1 integrin the addition of EGF resulted in significant increase of AP2-negative events (Figures 5F and 5G). These findings confirm that EGF selectively augments non-clathrin mediated endocytosis of β1 integrin.

Collectively, we conclude that EGF (and TNFα)-induced desialylation triggers Gal3 binding and clathrin-independent GL-Lect driven endocytosis of α3β1 integrin.

### EGF-induced desialylation is reversible

We previously hypothesized a link between GL-Lect driven endocytosis and retrograde trafficking to the Golgi apparatus where the glycosylation machinery including sialyltransferases is localized (Shafaq-Zadah et al., 2020). Using a HeLa cell line engineered for capture of retrograde cargoes (Johannes and Shafaq-Zadah, 2013) (Figure 6A), we observed that Gal3 itself is transported from the plasma membrane to the Golgi apparatus, and that this process was stimulated by incubation with EGF (Figure 6B).

**Figure 6.**
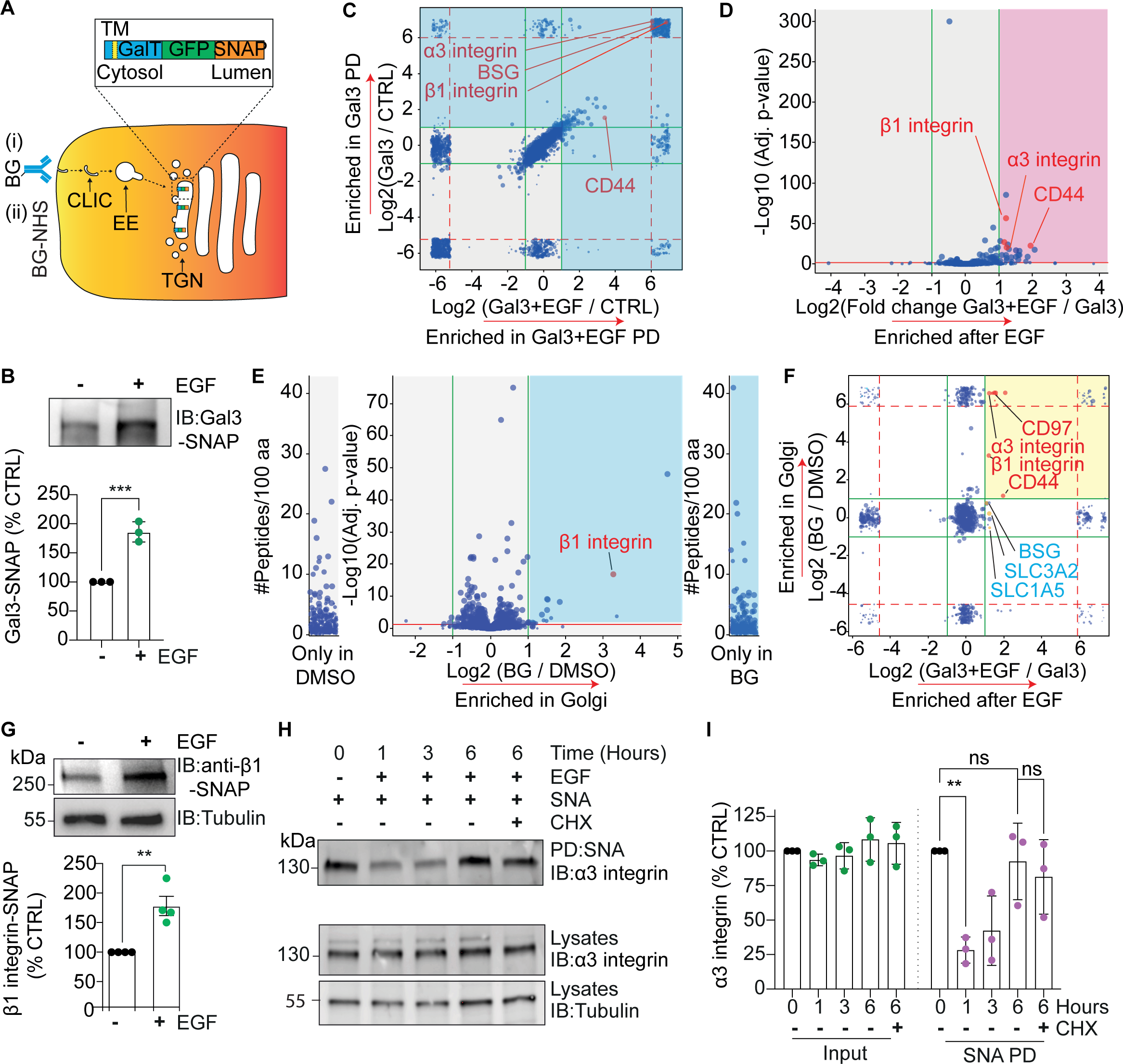
EGF-stimulated retrograde transport for resialylation. (A) Schematics of retrograde transport assay. Endocytic ligands (including antibodies against cell surface proteins) can either be directly coupled to benzylguanine (BG), as shown in (i), or the cell surface proteome can be modified with cell impermeable BG-NHS (ii). Proteins that undergo retrograde transport are captured in the Golgi by a covalent reaction with a GFP-tagged SNAPtag fusion protein that has been localized there. (B) HeLa cells stably expressing GalT-GFP-SNAP were continuously incubated for 4 h at 37 °C with BG-modified Gal3 before lysis and pulldown using GFP-trap beads. Western blot quantification from 3 independent experiments of Gal3-SNAP conjugate in pulldowns. Means ± SEM, unpaired t-test, *** p < 0.001. (C) Gal3 interaction proteomics. HeLa cells were incubated for 1 h on ice with Gal3-His in the presence or absence of EGF, followed by lysis, pulldown on cobalt-agarose beads, and quantitative mass spectrometry. Correlation plot of proteins. Blue quadrants indicate specific interactor of Gal3 in all conditions from 5 independent experiments. (D) Volcano blot of Gal3 interactors (blue quadrants from (C)) that are significantly (i.e., with 3 peptides and an adjusted p-value ≤ 0.05) enriched after EGF stimulation (red quadrant). (E) Retrograde proteomics. HeLa cells stably expressing GalT-GFP-SNAP were cell-surface modified on ice with NHS-PEG9-BG, followed by 16 h incubation at 37 °C, pulldown with GFP-trap beads, and quantitative mass spectrometry. High confidence retrograde cargoes are shown in the blue quadrant. (F) Comparison of Gal3 interactors and retrograde proteome. The yellow quadrant indicates high confidence Gal3 interactors that were enriched upon EGF stimulation and that also undergo retrograde transport. Note the presence of α3β1integrin and CD97 in this list. (G) 4 independent experiments as in (B) with BG-modified anti-β1 integrin antibodies. Quantification of anti-β1 integrin antibody-SNAP conjugates (IB anti-β1-SNAP) in GFP-trap pulldowns. Means ± SEM, unpaired t-test. (H) Resialylation analysis. MDA-MB-231 cells were incubated for 1 h on ice in the presence or absence of EGF, shifted in all conditions to 37 °C without EGF, and lysed at the indicated timepoints. ⍰3 integrin signal in SNA pulldowns (PD) was revealed by Western blotting. Note that ⍰3 integrin was still desialylated after 1 h of chase, and then became resialylated after 6 h of chase in a protein neosynthesis-independent manner (presence of cycloheximide (CHX). (I) Quantification of ⍰3 integrin signals from 3 independent experiments as in (H). Left, input; right, SNA PD. Means ± SEM, one-way ANOVA with Dunnett’s multiple comparison test, ns = p > 0.05, ** p < 0.01.

To gain a general view on the link between Gal3, EGF and retrograde transport, Gal3 binders at the plasma membrane were pulled down from EGF-treated or untreated Golgi SNAP-tag expressing HeLa cells and analyzed by quantitative mass spectrometry (Figure 6C). Of the Gal3 interactors that were present in either condition (blue quadrants from Figure 6C), 26 were significantly enriched after EGF stimulation (red quadrant in Figure 6D) in addition to 235 unique proteins (Table S3). Gene ontology revealed cell adhesion, migration and locomotion as some of the most highly enriched terms (Figure S3C), similarly to what was shown in Figure 2E for MDA-MB-231 cells. We then determined the retrograde proteome in these cells (Figure 6E) and used this to identify the EGF-stimulated Gal3 interactors, *i.e*., client proteins, that underwent retrograde transport to the Golgi (yellow quadrant in Figure 6F). Notably, α3β1 integrin, CD97 and a known Gal3 cargo, CD44 (Lakshminarayan et al., 2014), were amongst the hits (Tables S4 and S5). Using the antibody uptake protocol (Figure 6A), we confirmed that retrograde trafficking of the β1 and α3 integrin chains was indeed stimulated by EGF (Figures 6G and S3D).

We next tested if proteins acutely desialylated at the cell surface by EGF treatment undergo resialylation intracellularly. As shown above (Figures 2F and 2G), EGF treatment on ice led to a 36% reduction of α2-6-linked Sia modifications on α3 integrin. After 1 h of chase at 37 °C, during which receptor-bound EGF continues to signal, the reduction of α2-6 sialylation was as high as 72% (Figures 6H and 6I, 1 h lane). Strikingly, after 6 h of chase at 37 °C, the α2-6 sialylation of α3 integrin returned to pre-EGF treatment levels (Figures 6H and 6I, 6 h lane). Since this recovery was also observed when cells were chased in the presence of the protein biosynthesis inhibitor cycloheximide (Figure 6H and 6I, 6 h lane +CHX), we concluded that sialylation did not originate from newly synthesized α3 integrin.

Our data demonstrate the existence of an EGF-controlled reversible Sia glycan-based regulatory circuit, which through the EGF-induced desialylation and Gal3-dependent retrograde trafficking of plasma membrane glycoproteins provides access to the Golgi apparatus to reset their sialylation state.

### The stimulation of cell migration by EGF depends on desialylation

Endocytic trafficking is critical for complex cellular properties such as cell polarity and migration (Moreno-Layseca et al., 2019; Shafaq-Zadah et al., 2020; Sigismund et al., 2021). We previously reported that retrograde transport of β1 integrin from the cell surface to the Golgi apparatus is required for its polarized re-localization to the leading edge of migrating cells and thereby, for persistent cell migration (Shafaq-Zadah et al., 2016). Given that EGF is known to stimulate cell migration (Blay and Brown, 1985), we investigated whether this effect was dependent on the discovered regulatory de/resialylation circuit.

First, we studied the migration of MDA-MB-231 cells in cell-derived matrices (Kaukonen et al., 2017) obtained from telomerase immortalized fibroblasts. Velocity, accumulated distance, and notably Euclidean distance were increased by EGF (Figures 7A, 7B, S3E and S3F). Strikingly, this stimulatory effect of EGF was consistently abrogated in the presence of the Gal3 inhibitor I3 (Figures 7A, 7B, S3E and S3F), and by genetic or pharmacological interference with the EGFR tyrosine kinase, MEK, NHE1, Neu1 and Neu3 activities (Figures 7C, S3G and S3H).

**Figure 7.**
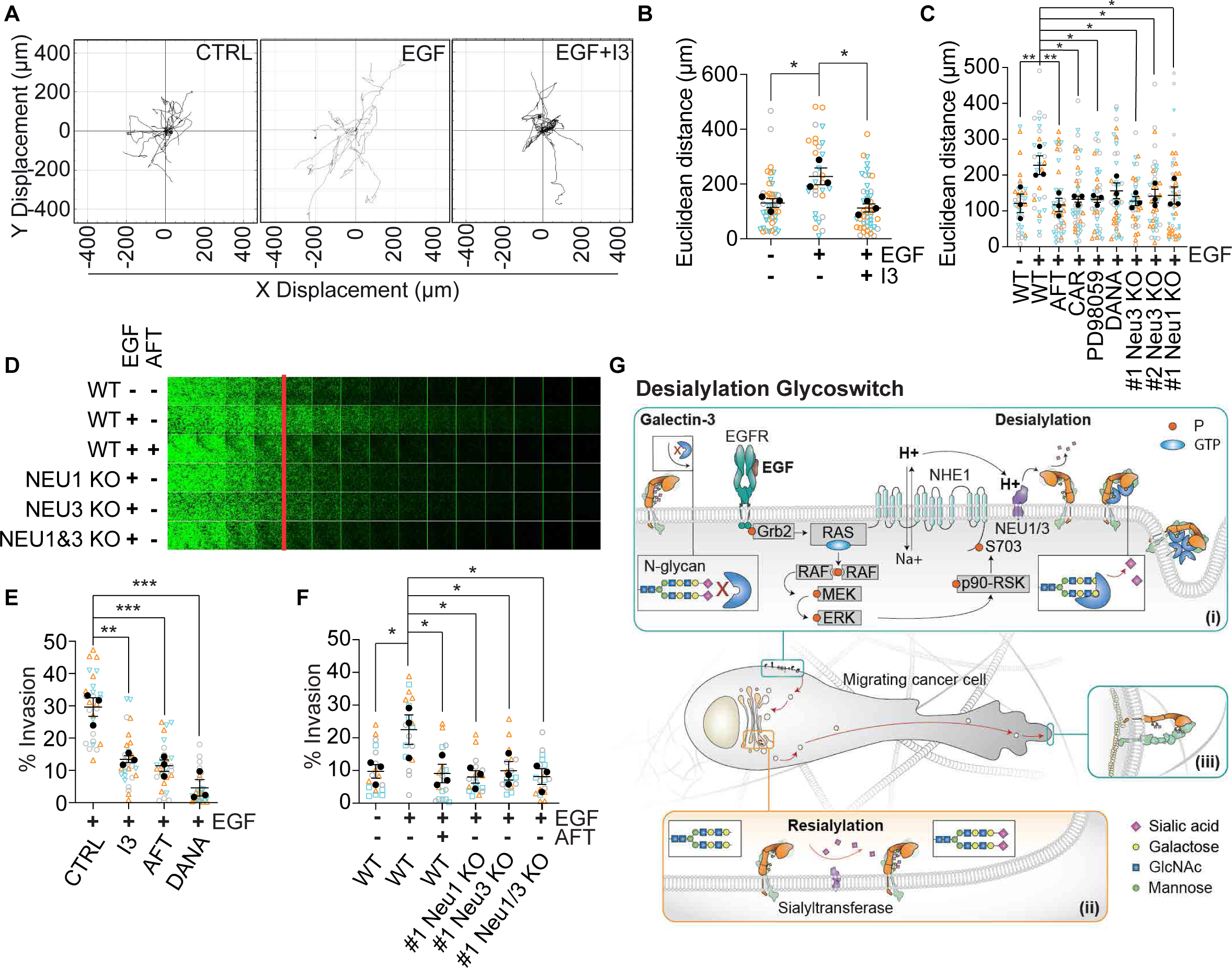
EGF-induced desialylation is required for invasive cell migration. (A) Cell migration. MDA-MB-231 cells were seeded under the indicated conditions on cell derived matrices and incubated for 16 h incubations at 37 °C in 1 % FCS. Individual tracks are represented. (B) The Euclidean distance of migration was determined as in (A) in 3 independent experiments in the absence (DMSO) or presence of the Gal3 inhibitor I3 and of EGF. (C) Experiment as in (B) in the absence (DMSO) or presence of the indicated drugs. (D) Inverted invasion assays. MDA-MB-231 cells were attached to fibronectin-spiked collagen plugs and allowed to invade against gravity towards an upper reservoir with or without EGF. Invasion was considered beyond the indicated red line. Knockout of the indicated neuraminidases inhibited invasion. (E-F) Quantification of invasion from 3 independent experiments as in (D) with the indicated inhibitors and KO cells. (G) Schematics depicting the desialylation glycoswitch model. See text for details. Means ± SEM are shown in this figure. Means from separate experiments are indicated by solid dots, and measurements of individual cells have different colored symbols for each experiment. One-way ANOVA with Dunnett’s multiple comparison test, * p ≤ 0.05, ** p < 0.01, *** p < 0.001.

Second, we measured the migration of MDA-MB-231 cells using an inverted invasion assay, where cells migrate through a 3D environment made from rat tail collagen supplemented with fibronectin, and in which EGF is used as a chemoattractant (Hennigan et al., 1994). As expected, the EGFR inhibitor afatinib (AFT) reduced cell migration to the level observed in the absence of EGF (Figures 7D and 7E). The same reduction in cell migration was found when *NEU1* and/or *NEU3* were genetically eliminated (Figures 7D and 7F), when general sialidase activity was inhibited with DANA, or Gal3 function with I3 (Figure 7E). Migration may have relied on α3β1 integrin which, despite being a major receptor for laminins, has also been reported to function with collagens and fibronectin (Chen et al., 2022; Elices et al., 1991). Alternatively, other α chains showed a similar response to EGF treatment as α3 integrin (Figure 2D), such as α2 integrin, a major collagen receptor (Staatz et al., 1990).

The stimulatory effect of EGF on cell migration clearly depends on all the outlined components of the EGF-induced de/resialylation circuit that we have discovered.

## Discussion

Based on our findings, we propose the existence of a regulatory process - termed desialylation glycoswitch - in which EGF triggers the immediate desialylation of glycans on select surface glycoproteins and their rapid internalization and retrograde transport to the Golgi apparatus (Figure 7G) for redesign of the glycan makeup and repurposing to accommodate new functions such as the dynamic and polarized localization to specialized areas of the plasma membrane. The desialylation glycoswitch represents a novel concept that challenges the perceived notion of the glycan makeup of plasma membrane proteins being static. Since most plasma membrane glycoproteins carry Sia capped glycans (Schjoldager et al., 2020; Varki, 2016), this mechanism has the potential to shape a vast array of biological functions in virtually all tissues.

In the current study, we focused on cell adhesion molecules, integrins, and their retrograde transport-dependent polarized localization to the leading edge of migratory cells (Shafaq-Zadah et al., 2016), which we show to be required for EGF-induced persistent cell migration. This signaling-induced dynamic localization to specialized areas of the plasma membrane may also operate in other cell types (Shafaq-Zadah et al., 2020), such as in activated T lymphocytes for the construction of immunological synapses (Carpier et al., 2018). Furthermore, we focused on EGFR as an inducer of desialylation in relation to Gal3-driven endocytic uptake, which revealed a novel facet of EGFR signaling. Other growth factors (e.g., TNFα), and other desialylation triggers may also operate the desialylation glycoswitch mechanism, such as depolarization in synaptic nerve terminals (Boll et al., 2020) and acute stress (Abe et al., 2019). The latter may function through α-adrenergic stimulation-induced increase of EGF levels in serum, which can become as high as 150 ng/mL (Byyny et al., 1974; Cohen and Elliott, 1963). Acidification of the extracellular milieux, as seen in cancer (Amith and Fliegel, 2013), may also sensitize tumor tissue for desialylation glycoswitch-mediated effects through the NHE1-dependent pH trigging mechanism that we uncovered in our study. It is furthermore notable that the maturation of neurons leads to an increase in Sia content (Govind et al., 2021).

EGF levels are found in the range of dozens to hundreds of ng/mL in body fluids such as saliva and urine (Byyny et al., 1974; Sigismund et al., 2005) (see Supplemental Materials for more details). Our finding that Gal3 binding to basal cells of the saliva-exposed tongue is decreased upon EGFR inhibition in living mice indicates that the desialylation glycoswitch may indeed operate *in vivo* in this tissue. A strong human genome-wide association study (GWAS) signal for estimated glomerular filtration rate traits are found in the identified desialylation glycoswitch component Neu1 (Wuttke et al., 2019), suggesting an involvement in protein resorption from urine by proximal tubule epithelial cells. Of note, the reduced Gal3 binding to cells from patients with genetically determined glycosylation disorders (Willems et al., 2016) in which N-glycan maturation or remodeling is deficient, including *NEU1* in type 1 sialidosis (Bonten et al., 1996; Sphranger et al., 1977), indicates that the desialylation glycoswitch mechanism may be disrupted, in part explaining reported clinical symptoms such as proteinuria and protuberant tongue (Ranganath et al., 2012). *In vivo* effects of glycan changes associated with *NEU1* may also be molecularly interpreted by other galectins, with overlapping and distinct specificities and functions (Johannes et al., 2018; Leffler and Barondes, 1986; Stowell et al., 2008; Tribulatti et al., 2012). Gal3 knockout mice have some distinct phenotypic deficiencies, but only mild effects in other cellular functions (Cao et al., 2002; Colnot et al., 1998; Hsu et al., 2000). This may be due to compensatory action by other galectins, or other mechanisms.

The EGF-induced trafficking cycle is poised for the repurposing of plasma membrane glycoproteins by changing their glycan makeup. Sialylation itself is an obvious candidate. A critical step is regulating the linkage (α2-3 or α2-6) of Sia capping of glycoproteins, as this has major consequences for recognition of glycan-binding proteins, such as the 15 members of the galectin family (Leffler and Barondes, 1986; Stowell et al., 2008; Zhuo and Bellis, 2011) (Gal3 in the current study), or Sia-binding immunoglobulin-type lectins (Siglecs) with their effects on inflammation and immune checkpoints (Läubli and Varki, 2020; Varki and Gagneux, 2012). We envision that Sia linkages can be tuned in function of pathophysiological situations following growth factor-induced desialylation at the plasma membrane and subsequent resialylation in the Golgi apparatus. Whether other types of glycosylation can also be edited through this cycle remains to be explored.

In conclusion, our desialylation glycoswitch concept elevates glycosylation to the level of dynamic regulatory processes. In separate study, we propose another mechanism, termed the conformational glycoswitch, by which the spatial rearrangement of N-glycans during conformational transitions of membrane glycoproteins triggers Gal3 oligomerization leading to GL-Lect driven endocytosis (Shafaq-Zadah et al., submitted). While the mechanistic wiring is different with covalent glycan modifications in the case of the desialylation glycoswitch and spatial glycan remodeling in the case of the conformational glycoswitch, both mechanisms are complementary and have in common the acutely triggered binding of galectins to glycans to drive endocytosis. Our findings of these glycoswitches, call for the reinterpretation of a plethora of biological interactions guided by glycans, for which dysregulation is causally linked to common human pathologies (Smith and Bertozzi, 2021; Stowell et al., 2015; Willems et al., 2016).

## Acknowledgments

We would like to thank Larry Fliegel (University of Alberta, Edmonton, Canada), Thomas Boltje and Christian Büll (Radboud University, Radboud Consortium for Glycoscience, Nijmegen, The Netherlands) for generously providing reagents and cell lines, Zhang Yang and Yoshiki Narimatsu (Copenhagen Center for Glycomics, Denmark) for advice and help on the generation of neuraminidase knockout CRISPR cells, Sara Sigismund (European Institute of Oncology, Milano, Italy) for helpful discussions on EGFR trafficking and signaling, Jean Salamero (Institut Curie, Paris, France), Thomas Van Zanten and Greeshma Pradeep (NCBS, Bangalore, India) for their support with the microscopy, and Jean Salamero for managing and providing funding for the Lattice Light Sheet Microscopy system. We would also like to acknowledge the CurieCoreTech facilities for Cytometry, Recombinant Proteins, Cell and Tissue Imaging (PICT-IBiSA), and the Nikon Imaging Centre at Institut Curie, member of the French National Research Infrastructure France-BioImaging (ANR10-INBS-01). The Johannes and SERPICO teams are members of Labex Cell(n)Scale (11-LABX-0038) and Idex Paris Sciences et Lettres (ANR-10-IDEX-0001-02 PSL). This wok was supported by grants from Mizutani Foundation for Glycosciences reference n° 200014 (LJ), Q-Life ANR-17-CONV-0005 (LJ), Agence Nationale pour la Recherche ANR-16-CE23-0005, ANR-19-CE13-0001-01, ANR-20-CE15-0009-01, ANR-22-CE11-0030-03 (LJ), Fondation pour la Recherche Médicale EQU202103012926 (LJ), Labex DCBiol ANR-11-LABX-0043 (LJ), ITMO Cancer 18CQ091 (LJ, SERPICO), France-BioImaging National Infrastructure ANR-10-INBS-04-07 (SERPICO), “La Région Île-de-France” n° EX061034 (DL), ITMO Cancer of Aviesan and INCa on funds administered by Inserm (N°21CQ016-00) for mass spectrometry (DL), Department of Atomic Energy, Government of India, under Project Identification No. RTI 4006, JC Bose fellowship from Department of Science and Technology (GoI) and Margadarshi Fellowship (IA/M/15/1/502018) (SM) and a Curie-NCBS Campus fellowship (EM) and ICMR PhD fellowship (RG), Novo Nordisk Foundation Grant NNF0067602 (TDM), The Novo Nordisk Foundation and Danish National Research Foundation (DNRF107) (HC), University Grants Commission (UGC) of India for graduate fellowship (RG), Cancer Research UK (DCRPGF\100002) (PC), European Union’s Horizon 2020 research and innovation programme under the Marie Skłodowska-Curie grant agreement No. 847718847718 (DG).

## Author Contributions

LJ, CW, EM, ML, HL for Conceptualization. LJ, CW, EM, AF, ED, M S-Z, TDM, RW, JH, CV-C, RG, SM for Methodology. EM, CW, AF, JH, CV-C, LL, ED, M S-Z, DB, TDM, DG, AP, YN for Investigation. EM, CW, AF, JH, CV-C, VC, AAS, BL, DL, ED, MS-Z for visualization. LJ, SM, RW, EM, PC Funding acquisition. LJ for Project administration. LJ, CW, SM, RW, PC for Project administration. LJ, EM, CW wrote the original draft. All authors edited and agreed to the final version of the manuscript.

## STAR Methods

### Cell lines and cell culture

All cell lines used in this study are from human origin unless otherwise stated. MDA-MB-231 (ATCC HTB-26), genome edited MDA-MB-231 (NEU1-/-, NEU3-/-, NHE1-/-), MDA-MB-468 NHE1-/-, MDA-MB-468 NHE1-/- rescued with NHE1-mEmerald, HN12, MEF (mouse), HeLa, SUM159-AP2-eGFP and NR6 (mouse) cells were grown in Dulbecco’s modified Eagle medium (DMEM), supplemented with 10% fetal bovine serum (FBS), 1 mM sodium pyruvate, 2 mM glutamine and penicillin/streptomycin (Thermo Fisher Scientific). Human fibroblast cultures from patients with genetic glycosylation defects were collected as part of clinical care. Residual, de-identified material was used in this study under ethics agreements from the Radboud University Medical Center, the Netherlands (2020-6588). The fibroblasts were cultured in M199 media (Sigma M2154), supplemented with 10% fetal bovine serum (FBS), 1 mM sodium pyruvate, 2 mM glutamine and penicillin/streptomycin (Thermo Fisher Scientific). All cell lines were regularly tested for mycoplasma contamination.

### Growth factors

Human growth factors were purchased from Miltenyi Biotech: EGF (130-097-751), FGF-1 (130-095-790), FGF-2 (130-104-918), PDGF-BB (130-108-163), TNF-alpha (130-094-015), VEGF (130-109-395), and from Novus Biologicals: human TGF-alpha (NBP2-3643).

Unless stated otherwise, EGF was used at the saturating concentrations of 100 ng/mL in the experiments of this study to achieve maximal robustness throughout the multitude of experimental approaches that were used. This concentration is within the physiological range of several body fluids such as serum under stress conditions (Byyny et al., 1974; Cohen and Elliott, 1963), mouse saliva, urine, milk (Beardmore and Richards, 1983; Byyny et al., 1974; Hayashi and Sakamoto, 1988), bile (Grau et al., 1994), and prostate fluid (Gann et al., 1997). As shown in Fig. 1C, Gal3 binding to cells is stimulated by as little as 1 ng/mL, however. We envision that the desialylation glycoswitch is triggered locally in the environment of specific tissues, and possibly also systemically in acute situations such as stress.

### Antibodies

Anti-⍰3 integrin (MA5-28565, ASC-1, Thermo Fisher Scientific, immunofluorescence and antibody uptake), anti-⍰3 integrin (66070, Proteintech, for Western blotting), anti-β1 integrin (NBP2-52708, Novus, clone K20, immunofluorescence and antibody uptake), anti-pEGFR Y1068 (3777, Cell Signaling Technology), anti-P-p90RSK (Ser380) rabbit monoclonal antibody D3H11 (11989S, Cell Signaling Technology), anti-Phospho-p44/42 MAPK (Erk1/2) (Thr202/Tyr204) (9101, Cell Signaling Technology), anti-p44/42 MAPK (Erk1/2) (9102, Cell Signaling Technology); anti-actin (A5316, Sigma), anti-tubulin (T5168, Sigma), anti-SNAP (New England Biolabs, P9310S), anti-EGFR (D38B1, Abcam), anti-CD97 (clone VIM3b, Biolegend).

### Antibody modification

Antibodies were coupled for 16 h at 16 °C with the NHS-reactive versions of AcidiFluor Orange560 (GC302, Goryo Chemical), Atto647N (AD 647N, ATTO-TEC), benzylguanine (S9151S, NEB), or Cy3 (PA23001, Cytiva), or aldehyde-activated HRP (31487, Thermo Fisher Scientific) using a molar excess of 6 in 125 mM sodiumbicarbonate at pH 8 and 6% glycerol. The coupling product was purified by size exclusion chromatography (Zeba spin 40K, 87767, Thermo Fisher Scientific) or for HRP SD75 3.2/300 FPLC column (Cytiva) in PBS (for HRP). Labeling efficiency was determined by spectrometric analysis on NanoDrop (Thermo Fisher Scientific).

### Lectins

Sambucus Nigra lectin (SNA) versions: SNA-Biotin (B-1305-2, Vector labs), SNA-Fluorescein (FL-1301-2, Vector labs), SiaFind^TM^ α2,3-Specific lectin, biotinylated (SK2301B, Lectenz), SiaFind^TM^ Pan-Specific lectin, biotinylated (SK0501B, Lectenz).

### Inhibitors

N-Acetyl-2,3-dehydro-2-deoxyneuraminic acid (DANA) (D9050, Sigma, working concentration: 1 mM), RSK Inhibitor II (CAS 501437-28-1, Calbiochem, working concentration: 200 nM), PI-103 (CAS 371935-74-9, Calbiochem, working concentration: 100 nM), PD 98059 (CAS 167869-21-8, Calbiochem, working concentration: 10 µM), Cariporide (CAR) (SML1360, Sigma, working concentration: 200 nM), afatinib (AFT) (SYN-1100, Euromedex, working concentration: 100 nM), Gal3 inhibitor I3 (working concentration: 10 µM), sialostatin (STI) (also called, SiaFNEtoc, Thomas Boltje, Radboud-University, working concentration 10 µM, treatment for 72 h), cycloheximide (239763-Sigma, working concentration: 100 µg/ml). For all experiments, cells were pre-treated with inhibitors in serum-free media for 30 min, unless stated otherwise.

### Other reagents

Transferrin Alexa Fluor546 (T23364, Thermo Fisher Scientific), complete mini EDTA-free Protease Inhibitor Cocktail (11836170001, Roche), Phosphatase Inhibitor Cocktail 1 (Sigma # P2850).

### siRNA-mediated knockdown

For knockdown experiments, cells were seeded at 30% confluency and grown for 24 h. siRNA transfection was performed with HiPerfect (Qiagen) according to manufacturer’s instructions, using 5 µL HiPerfect and 25 nM siRNA final for 6-well plates. Knockdown was validated by Western blotting after 72 h. See Table S6 for siRNA sequences.

### Recombinant proteins

C-term His-tagged Gal3 (Gal3-His) and Cys-Gal3-His (pHisParallel2) were used as described in (Lakshminarayan et al., 2014). Cys-Gal3-His was generated using the corresponding primers of Table S6. His-tagged Gal3 proteins were expressed overnight at 20 °C in Rossetta2-pLysS strains (Novagen) using LB media, supplemented with 60 μM IPTG. 95% purity as measured by Coomassie staining was achieved using Cobalt-resin (Pierce) affinity and gel filtration (Superdex75 16×600) chromatography. Purification buffers were PBS pH 7.3, or 20 mM HEPES pH 7.3, 150 mM NaCl supplemented with 10 mM lactose and 0.2 mM TCEP (20 μM TCEP during FPLC purification) if subsequent coupling with HRP was performed. For Alexa Fluor488 labeling, Gal3 (2 mg/mL) in PBS was mixed with amine-reactive (5-TFP ester) Alexa488 (Thermo Fisher Scientific) in a molar ratio of 1:4 and incubated for 16 h at 16 °C with 500 rpm agitation. The mixture was purified using PD-10 columns (Sigma), snap-frozen, and stored at −80 °C. Labeling efficiencies were between 1.2 and 1.8 by spectrophotometric analysis. Maleimide-activated horseradish peroxidase (HRP; Thermo Fisher Scientific) coupling to Cys-Gal3-His was performed for 12 h at 4 °C in 20 mM HEPES pH 7.3, 150 mM NaCl, 10 mM lactose, 20 μM TCEP at a molar ratio of 1:2. For purification, gel filtration chromatography was performed using a Superdex75 10/300 column. HRP-Gal3-His in 50% glycerol was snap-frozen and stored at −80 °C.

### Gal3 binding assays on cells in culture

#### Exogenous Gal3

When indicated, cells on coverslips were serum starved by incubation for 1 h (growth factor screen and NEU1/3 knockout MDA-MB-231 cells) or 30 min (all other experiments unless otherwise stated) at 37 °C in serum-free DMEM in the presence or absence of indicated inhibitors, incubated for 1 h on ice with 200 nM of Alexa Fluor-488 labeled Gal3 in serum-free DMEM, washed 3x in ice cold PBS, fixed in 4% PFA/PBS and quenched in 50 mM NH_4_Cl in PBS. For pH-buffer assays, cells were serum-starved for 1 h, washed in DMEM, and incubated for 1 h on ice in 25 mM HEPES pH 7.4 or pH 8.4 buffers in the presence or absence of 100 ng/mL EGF. In the pH rescue condition, cells were washed and incubated for 1 additional h on ice in pH 7.4 buffer in the presence of 100 ng/ml EGF. Cells were fixed in 4% PFA/PBS and quenched in 50 mM NH_4_Cl in PBS, before incubation in DMEM pH 7.4, and 200 nM Alexa-488 Gal3. For the pH 5.5 assay, cells were incubated for 30 min at 37 °C in serum-free DMEM in the presence or absence of 1 mM DANA before incubation for 1 h on ice in PBS pH 5.5 or PBS pH 7.4 in the presence or absence of 100 ng/ml EGF and 1 mM DANA. Cells were then washed and further incubated for 1 h on ice in serum free DMEM with 200 nM Alexa488-Gal3 in DMEM and in the presence or absence of 1mM DANA. Cells were fixed in 4% PFA/PBS, quenched in 50 mM NH_4_Cl in PBS, and mounted using Fluoromount-G. Images were recorded using confocal microscopy on an inverted Eclipse Ti-E (Nikon) or spinning disk CSU-X1 microscope (Yokogawa) with integrated Metamorph software by Gataca Systems, 60x CFI Plan Apo Lambda, in stacks of 16 images with a depth of 0.2 μm each. Images were processed in ImageJ. Individual cells were analyzed on SUM stacks after background removal. Values were normalized separately to control, and outliers were identified with ROUT analysis where appropriate. Mean fluorescent intensity was calculated for individual cells and values expressed as a % of the mean of the control condition.

#### Endogenous Gal3

MDA-MB-231 cells were cultured for two days in DMEM 10% FCS on coverslips. On the second day, media was removed and kept on ice (preconditioned media contains secreted Gal3). Cells were washed twice and incubated for 30 min at 37 °C in serum free DMEM. The pre-conditioned media contained PBS vehicle or 100 ng/mL EGF. Cells were cooled down on ice for 5 min and starvation media was exchanged with ice-cold preconditioned media (+/− EGF), followed by incubation for 1 h on ice. Unbound proteins were removed by two washes with ice-cold PBS^++^. Cells were fixed for 20 min at room temperature with 4% PFA (Electron Microscopy Sciences), quenched for 10 min with 50 mM NH_4_Cl, blocked for 30 min with 0.3% BSA (IgG-free, Jackson Immuno Research) and 0.4% cold-water fish-gelatine (Aurion) in PBS, and labeled with 1 µg/mL anti-Gal3 primary antibody (gift of Fu-Tong Liu, Institute of Biomedical Sciences, Academia Sinica, Taipei, Taiwan) and secondary antibody (anti-goat Alexa Fluor 488, Jackson Immuno Research, 1:400). Z-stacks of samples were acquired by confocal microscopy (Leica SP8) using a 63x oil immersion objective. For quantification, images were background subtracted, cellular regions were marked, and the total fluorescence intensity of the merged z-stack was measured in these regions using ImageJ (NIH). Intensities were normalized separately to the untreated control sample.

### Western blotting

For signaling experiments, MDA-MB-231 cells were lysed for 10 min on ice with gentle rocking in RIPA buffer (150 mM NaCl, 1% NP-40, 0.1% SDS, 50 mM Tris pH 7.4), 2 mM sodium orthovanadate and phosphatase inhibitor cocktail 1 (Sigma, # P2850), or for patient fibroblasts cells in 0.5% NP-40, 0.5% Triton X-100 in PBS. Lysates were cleared at 16,000 x g for 10 min at 4 °C in a bench top centrifuge. BCA assay was used to determine protein concentration, equal amounts of protein (typically 10-25 µg) were loaded onto 4-20% gradient or 7.5% Tris-Glycine precast gels (Biorad), and transferred in Towbin’s buffer (25 mM Tris pH 8.3, 192 mM glycine, 20% methanol) onto 0.2 µm pore nitrocelluose membrane (Amersham Protran #10600001), using the Trans-blot (Bio-rad) semi-dry transfer system. Antibody incubations were performed overnight at 4 °C in 5% BSA-TBS-T, unless otherwise stated. Quantification of bands was performed using Image J and normalized to loading control.

### Antibody uptake

Cells were serum starved for 30 min at 37 °C before continuous incubation for 10 min at 37 °C with 1 µg/mL of the indicated antibodies or 5 μg/mL Alexa488-transferrin in DMEM supplemented with 0.2% fatty acid free BSA. Cells were placed on ice, cell surface accessible antibodies or transferrin were removed by 3 washes in 0.5 M glycine pH 2.2, and cells were then fixed in 4% PFA/PBS and quenched in 50 mM NH_4_Cl in PBS. Images were captured using confocal microscopy on an inverted Eclipse Ti-E (Nikon) or spinning disk CSU-X1 (Yokogawa) microscopes with integrated Metamorph software by Gataca Systems, 60x CFI Plan Apo Lamda. Stacks of 16 images with a depth of 0.2 μm depth were acquired. Images were processed in ImageJ, and individual cells were analyzed on SUM stacks after background removal. Mean fluorescent intensity was calculated for individual cells and values expressed as a % of the mean of the control condition.

### pH imaging

MDA-MB-231 cells were seeded in imaging dishes. Cells were incubated for 15 min at 37 °C with 200 nM cariporide (CAR) or DMSO, followed by an incubation with 1 μg/mL anti-⍰3 integrin-AcidiFluor Orange for 4 min. Cells were washed in imaging media (150 mM NaCl, 5 mM KCl, 1.2 mM CaCl_2_, 2.2 mM MgCl_2_, 20 mM HEPES pH 7.3), and imaged for 10 min at 37 °C in the presence of absence of 100 ng/mL EGF sequentially in TIRF and epifluorescence modes with an interval of 3 s per frame. For the pH test, cells were prepared as above and imaged in 200 μl pH 7.4 buffer before addition of 10 volumes of pH 4 buffer (150 mM NaCl, 20 mM sodium acetate) at the indicated timepoints. TIRF images were analyzed in ImageJ track mate to identify spots. Tracks that persisted over 3 frames were quantified as total number of tracks and the peak fluorescence intensities were extracted for each individual track. ROUT analysis was used to identify outliers. Image sequences are walking averages of 3 consecutive frames generated using ImageJ. Histograms were generated from using Image J to measure the fluorescence values in each channel in a defined area indicated by arrows. Images were captured on Nikon eclipse Ti microscope with an Agilent MLC100 combiner with 543 nm laser line, 60x CFI Plan Apo Lambda, CMOS camera.

### Retrograde transport with purified benzylguanine-coupled ligands

24 h before experiments, HeLa cells stably expressing Golgi-resident GalT-GFP-SNAP protein were seeded on 6-well plates in complete DMEM medium to reach 80% confluency the day of experiment. Cells were first serum-starved for 30 min at 37°C, followed by continuous incubation for 4 h at 37 °C in the presence or absence of 100 ng/mL EGF with 10 μg/mL of benzylguanine-coupled anti-β1 integrin antibody or with anti-α3 integrin antibody, or with 5 μg/mL of benzylguanine-coupled Gal3 in in serum-free DMEM. Unreacted GalT-GFP-SNAP was quenched by incubating cells for 30 min at 37 °C with 10 μM SNAP-Cell-Block (NEB) diluted in serum-free DMEM. After 3 washes with PBS^++^ (PBS, 0.5 mM MgCl_2_, 1 mM CaCl_2_), cells were lysed for 30 min on ice with 0.5 ml TNE (10 mM Tris pH 7.4, 150 mM NaCl, 5 mM EDTA) supplemented with 1% NP-40 and protease inhibitor cocktail (Sigma Aldrich, Ref. P8849), with regular spreading of the lysis solution. Cells were then scraped off and lysates centrifuged at 16,000 x g for 15 min at 4 °C. Supernatants were incubated with 40 μL bed volume of protein G sepharose beads for overnight pulldown at 4 °C on a rotating wheel. Beads were washed 3 times with TNE 0.1% NP-40. Proteins were eluted with 40 μL of 1.5x SDS-sample buffer and denaturated for 10 min at 95 °C. Pulldown samples were loaded on SDS-PAGE and analyzed by Western blotting with anti-SNAP antibody. Total lysate samples were analyzed by Western blotting using anti-tubulin antibody for SNAP signal normalization.

### Biological approaches leading to mass spectrometry

#### SNA pulldown on MDA-MB-231 cells

Cells were treated with STI for 72 h before incubation for 1 h on ice with 5 μg/mL SNA-biotin in the presence and absence of 100 ng/mL of EGF, washed with ice-cold PBS, and lysed in 1% NP-40/PBS containing protease inhibitor cocktail. Lysates were passed 10 times through a 23G needle and cleared by centrifugation at 16,000 x g for 10 min at 4 °C. 1 mg of protein lysate, as determined by the BCA assay, was incubated for 1 h at 4 °C under rotation with M280-Strepavdin-dynabeads. Beads were washed 3 times in 1% NP-40/PBS, 3 times in 0.1% NP-40/PBS, followed by their resuspension in sample buffer for Western blotting. Alternatively, beads were washed 3 times in 25 mM ammonium bicarbonate for mass spectrometry.

#### Gal3 pulldown on MDA-MB-231 cells

Cells were seeded for 24 h at 37 °C in 10 cm dishes to reach a final density of 60%, washed with serum-free DMEM, starved for 30 min at 37 °C, and subsequently incubated for 1 h at 4 °C in serum-free media containing 5.2 μg/mL of Gal3–His. After rinsing twice with PBS, cells were scraped in 1 mL lysis buffer (PBS, 0.5% Triton X-100, 0.5% NP-40, protease inhibitor cocktail). Lysates were kept rocking on ice for 10 min and cleared (16,000 x g, 10 min, 4 °C). The supernatant was incubated for 1 h at 4 °C with HisTag Dynabeads (Thermo Fisher Scientific). Beads were washed twice with lysis buffer (without protease inhibitors), once with 150 mM ammonium bicarbonate 0.1% NP-40 buffer and processed for mass spectrometry.

#### Gal3 pulldown on HeLa cells

Cells were seeded for 24 h at 37 °C in 10 cm dishes to reach a final cell density of 60%. Cells were washed with serum-free DMEM and starved for 30 min at 37 °C. After one wash with PBS++ (PBS, 0.5 mM MgCl2, 1 mM CaCl2), cells were incubated for 10 min at 4 °C in lactose buffer (20 mM HEPES pH 7.3, 200 mM lactose, 45 mM NaCl, 5 mM KCl, 1 mM MgCl_2_, 1 mM CaCl_2_) to remove endogenous, cell surface bound Gal3. Cells were extensively washed with PBS++ (3 times with 20 mL), and subsequently incubated for 45 min at 4 °C in PBS++ (negative control) or in PBS++ containing 4 µg/mL of Gal3–His. After rinsing twice with PBS, cells were scraped in 0.8 mL lysis buffer (PBS, 0.5% Triton X-100, 0.5% NP-40, protease inhibitor cocktail). Lysates were kept rocking on ice for 30 min and cleared (16,000 x g, 10 min, 4 °C). Supernatants were incubated for 1 h at 4 °C with HisPur-Cobalt-Agarose (Thermo Fisher Scientific), washed twice with lysis buffer (without protease inhibitors), and once with 150 mM ammonium bicarbonate 0.1% NP-40 buffer. For protein deglycosylation, the 30 μL bead suspension was supplemented with 500 Units of glycerol-free PNGase F (New England Biolabs), incubated for 2 h at 37 °C, diluted with 50 μL 0.1% NP-40, inactivated for 10 min at 75 °C, and processed for mass spectrometry.

#### Retrograde transport

24 h before experiment, HeLa cells stably expressing Golgi-resident GalT-GFP-SNAP protein were seeded on 10 cm dishes in complete DMEM medium to reach 60% confluency the day of experiment. Cells were chilled on ice for 10 min, washed 3 times with cold PBS supplemented with 0.5 mM CaCl2 and 1 mM MgCl2 (PBS++). Cell surface modification was performed by incubating cells for 1 h at 4 °C with 1 mM NHS-PEG9-BG (Johannes and Shafaq-Zadah, 2013), diluted in 1 ml cold PBS++ just prior to the experiment, with regular spreading of the solution. Cells were then washed 3 times with PBS++ and further incubated overnight at 37 °C, 5% CO2. To avoid post-lytic benzylguanine/SNAP reactions, unreacted GalT-GFP-SNAP was quenched by incubating cells for 30 min at 37 °C with 10 μM SNAP-Cell-Block (NEB) diluted in complete DMEM. After 3 washes with PBS++, cells were lysed for 30 min at 4 °C with 1 mL TNE (10 mM Tris, 150 mM NaCl, 5 mM EDTA), supplemented with 1% NP-40 and proteases inhibitor cocktail (lysis buffer), with regular spreading of lysis solution. Cells were then scraped, and lysates centrifuged at 16,000 x g for 15 min at 4 °C. Post-nuclear supernatants were incubated overnight at 4 °C on a rotating wheel with 30 μL bed volume of GFP-Trap MA beads (Proteintech). Beads were washed 3 times with TNE washing buffer (10 mM Tris, 150 mM NaCl, 5 mM EDTA and 0.1% NP-40). After the third wash, beads were resuspended in 30 µL of lysis buffer. For protein deglycosylation, the 30 μL bead suspension was supplemented with 500 Units of glycerol-free PNGase F (NEB), incubated for 2 h at 37 °C, diluted with 50 μL 0.1% NP-40, inactivated for 10 min at 75 °C, and processed for mass spectrometry.

#### Sample preparation

Five biological replicates were processed. For each sample, 100 µL bead suspension in 25 mM ammonium bicarbonate was digested for 1 h at 37 °C by adding 0.2 µg of trypsine/LysC (Promega). Samples were desalted in custom-made C18 StageTips packed by stacking one AttractSPE® disk (#SPE-Disks-Bio-C18-100.47.20 Affinisep) and 2 mg beads (#186004521 SepPak C18 Cartridge Waters) into a 200 µL micropipette tip for all samples, except retrograde transport samples which were first purified to remove detergents in tips containing SDB-RPS (AttractSPE Disk Bio SDB-RPS, Affinisep). Peptides were eluted using a ratio of 40:60 MeCN:H2O, 0.1% formic acid, and vacuum concentrated to dryness. Peptides were reconstituted in injection buffer (0.3% TFA) before liquid chromatography tandem mass spectrometry (LC-MS/MS) analysis.

### Mass spectrometry

#### LC-MS/MS analysis

Peptides from SNA pulldowns on MDA-MB-231 cells were separated by reversed phase liquid chromatography (LC) on an RSLCnano system (Ultimate 3000, Thermo Fisher Scientific) coupled online to an Orbitrap Fusion Tribrid mass spectrometer (Thermo Fisher Scientific). Peptides were trapped in a C18 column (75□μm inner diameter□×□2□cm; nanoViper Acclaim PepMap^TM^ 100, Thermo Fisher Scientific) with buffer A (2:98 MeCN:H2O in 0.1% formic acid) at a flow rate of 3.0□µL;/min over 4□min. Separation was performed using a 40□cm□×□75□μm C18 column (Reprosil C18, 1.9□μm, 120□Å, Pepsep PN : PSC-40-75-1.9-UHP-nC), regulated to a temperature of 40□°C with a linear gradient of 3% to 32% buffer B (100% MeCN in 0.1% formic acid) at a flow rate of 150□nL/min over 91□min. MS1 data were collected in the Orbitrap in range m/z 350-1550 (120,000 resolution; maximum injection time (IT) 100 ms; AGC 4×10E5). Charge states from 2+ to 6+ were required for MS2 analysis, and a 30 s dynamic exclusion window was used. MS2 scan were performed in the ion trap in rapid mode with HCD fragmentation (isolation window 1.6 Da; NCE 28%, maximum IT 35 ms; AGC 10E4).

For Gal3 pulldown on MDA-MB-231 cells and retrograde transport samples, LC was performed with an RSLCnano systeme coupled online to an Orbitrap Exploris 480 mass spectrometer (Thermo Fisher Scientific). Peptides were trapped onto the C18 column with buffer A at a flow rate of 3 µL/min or 2.5 µL/min respectively over 4 min. Separation was performed on a 50 cm nanoviper column (i.d. 75 µm, C18, Acclaim PepMap^TM^ RSLC, 2 μm, 100Å, Thermo Scientific) regulated to a temperature of 40 °C or 50 °C with a linear gradient of 3% to 29% or 2% to 30% buffer B respectively at a flow rate of 300 nL/min over 91 min. MS full scans were performed in the ultrahigh-field Orbitrap mass analyzer in ranges m/z 375–1500 (resolution of 120□000 at m/z 200; maximum IT 25 ms; AGC 300%). The top 20 most intense ions were subjected to Orbitrap for further fragmentation via high-energy collision dissociation (HCD) activation and a resolution of 15□000 with the auto gain control (AGC) target set to 100%. We selected ions with charge states from 2+ to 6+ for screening. Normalized collision energy (NCE) was set at 30 and the dynamic exclusion of 40 s.

For Gal3 pulldown on HeLa cells, LC was performed as above with an RSLCnano system coupled online to an Orbitrap Eclipse mass spectrometer (Thermo Scientific). Peptides were trapped on a C18 column with buffer A at a flow rate of 3.0 µL/min over 4 min. Separation was performed at a temperature of 50 °C with a linear gradient of 2% to 25% buffer B at a flow rate of 300 nL/min over 91 min. MS1 data were collected in the Orbitrap in ranges m/z 375–1500 (120,000 resolution; maximum injection time 60 ms; AGC 4 x 10E5). Charges states between 2+ and 7+ were required for MS2 analysis, and a 60 s dynamic exclusion window was used. MS2 scans were performed in the ion trap in rapid mode with HCD fragmentation (isolation window 1.2 Da; NCE 30%; maximum IT 35 ms; AGC 10E4).

#### Data analysis

For identification, the data were searched against the Homo sapiens (UP000005640_9606) UniProt database using Sequest HT through Proteome Discoverer (v.2.2 or v.2.4). Enzyme specificity was set to trypsin and a maximum of two-missed cleavage sites were allowed. Oxidized methionine, N-terminal acetylation, methionine loss and methionine acetylation loss were set as variable modifications. Asn to Asp (N) modification was also set as variable modification for sample with PNGase F incubation. Maximum allowed mass deviation was set to 10□ppm for monoisotopic precursor ions. For fragment ions, it was set respectively to 0.02 Da and 0.6□Da for the Orbitrap Exploris and the Orbitrap Fusion & Eclipse data. The resulting files were further processed using myProMS (Poullet et al., 2007) v.3.9.3 (https://github.com/bioinfo-pf-curie/myproms). False-discovery rate (FDR) was calculated using Percolator (The et al., 2016) and was set to 1% at the peptide level for the whole study. The label-free quantification was performed by peptide extracted ion chromatograms (XICs), reextracted by conditions and computed with MassChroQ (Valot et al., 2011) v.2.2.21.

For protein quantification, XICs from proteotypic peptides shared between compared conditions (TopN matching) with missed cleavages were used. Normalization at the peptide level was applied on the total signal to correct the XICs using Median and scale correction, or without when using only the restricted protein list of significantly upregulated proteins compared to the control. Only Figure 2D was normalized to amount of initial material. To estimate the significance of the change in protein abundance, a linear model (adjusted on peptides and biological replicates) was performed, and p-values were adjusted using the Benjamini–Hochberg FDR procedure. In order to eliminate non-specifically isolated proteins, only proteins with at least three total peptides in all replicates (n = 5), a two-fold enrichment, an adjusted p-value ≤ 0.05, and for unique proteins three total peptides, unless otherwise stated. Proteins selected with these criteria were further used for quantitative data analysis.

#### Quantitative data analysis

To identify SNA binders responsive to EGF, a list of 93 proteins was first generated from proteins meeting the aforementioned criteria in Gal3 vs CTRL pulldown and for SNA vs SNA+STI. Proteins found in this list were then analyzed for their loss in SNA pulldown after EGF stimulation by plotting on a volcano graph and generating a list of proteins with a p-value ≤ 0.05 and exceptionally here a cut off of >1.35-fold enrichment. To identify Gal3 interactors responsive to EGF, we generated a list of proteins meeting the aforementioned criteria for Gal3 vs CTRL and Gal3+EGF vs control. These proteins were then plotted on a Volcano plot for Gal3+EGF vs Gal3 pulldowns, and a list of EGF sensitive Gal3 interactors was thereby generated for proteins that had two-fold enrichment and p value < 0.05. For the cross-reference of the retrograde proteome and Gal3 interactors, we plotted benzylguanine vs CTRL and Gal3+EGF vs Gal3 on a correlative plot. Proteins that were common to both data sets were highlighted. All lists were generated using myProMS.

#### Gene ontology (GO) analysis

GO functional enrichment analysis was performed using myProMS by using the complete ontology file and the subsequent list of “biological processes” terms were summarized using Revigo software using SimRel Cluster analysis methodology(Supek et al., 2011), the resulting graphs were generated using R. For each data point the size of the spot equals -LOG2 p-value for each GO term, spot color equals - LOG10 number of annotations that have been summarized under each category. The mass spectrometry proteomics data have been deposited for all pulldowns and subsequent MS raw datasets to the ProteomeXchange Consortium via the PRIDE partner repository (Perez-Riverol et al., 2022) with the dataset identifier PXD041450. Reviewer account detail are: Username, project accession PXD041450, password YuWk8W9e.

### Resialylation assay

MDA-MB-231 cells were incubated for 1 h on ice in the presence or absence of 100 ng/mL EGF, after which cells were incubated at 37 °C in full media +/− cyclohexamide (100 µg/ml), and lysed at the indicated timepoints in 0.5% Triton X-100, 0.5% NP-40 in PBS buffer containing EDTA free protease inhibitors. 750 µg of lysate, determined by BCA assay, was incubated for 1 h at 4 °C under rotation with 5 µg/mL SNA-biotin and M280-Strepavdin-dynabeads. Beads were washed 3 times in 1% NP-40/PBS and resuspended in sample buffer. Samples and 1% of input were run on 7.5% Tris-Glycine precast gels (Biorad) and processed for Western blotting.

### Immunoprecipitation of **⍰**3 integrin

MDA-MB-231 cells were serum starved for 1 h at 37 °C, incubated for 1 h on ice with 1 µg/mL of anti-⍰3 integrin (ASC-1) in 0.2% fatty acid free BSA/DMEM, washed 3 times in PBS, incubated for 1 h on ice in the presence of absence of 100 ng/mL EGF washed 3 times with PBS, and lysed in 2.5% Elugent, 300 mM NaCl. 400 µg of lysate, determined by BCA assay, were incubated for 1 h at 4 °C with protein A/G beads, which were washed 3 times in lysis buffer. Samples were analyzed on stain-free gels, then submitted to Western blotting and probed with SNA-biotin, according to the manufacturer’s instructions.

### Genome editing

MDA-MB-231 CRISPR KO cell lines for Neu1 or Neu3 were generated following the published protocol for genome editing using the Indel Detection by Amplicon Analysis (IDAA) (Lonowski et al., 2017). MDA-MB-231 cells were co-transfected with the specific gRNA pU6-plasmid (Table S6) and the Cas9-GFP pBKS vector (#68371, Addgene) using Ingenio electroporation solution (Mirus). 24 h after transfection, cells were FACS sorted into 96-well plates for GFP expressing cells. 96-wells were duplicated. One set was used for IDAA analysis and subsequent sequencing to validate the knockout. The second set was used for generating a permanent knockout cell line stock and tested for mycoplasma contamination.

### Migration on cell-derived matrices

Cell-derived matrices were generated according to the method developed by the Yamada team (Cukierman et al., 2001). Briefly, 6/12-well plastic plates (Corning) were coated with 0.2% gelatin (v/v, Sigma Aldrich), crosslinked with 1% glutaraldehyde (v/v, Sigma Aldrich) and quenched with 1 M glycine (Thermo Fisher) before telomerase immortalised fibroblasts (TIFs) were confluently seeded. DMEM medium supplemented with 0.25% ascorbic acid (v/v, Sigma Aldrich) was changed every 48 h for 8 days. Cells were denuded with extraction buffer (20 mM ammonium hydroxide (NH_4_OH); 0.5% (v/v) Triton X-100) to leave only matrix. Finally, DNAse was used for cleavage of phosphodiester linkages in the DNA backbone. Wildtype, Neu1, or Neu3 knockout MDA-MB-231 cells were then seeded sparsely in these cell-derived matrices, left to spread for 4 h at 37 °C, pretreated or not as indicated with I3 / afatinib (AFT) / cariporide (CAR) / PD98059 / DANA / DMSO vehicle control for 1 h and with EGF for 15 min at 37 °C prior to imaging cell migration. Cells were incubated at 5% CO_2_ for 16 h at 37 °C in DMEM with 1% FCS, and recorded using an Eclipse Ti inverted microscope (Nikon) with a 20x / 0.45 SPlan Fluor objective and the Nikon filter sets for brightfield and a pE-300 LED (CoolLED) fluorescent light source with imaging software NIS Elements AR.46.00.0. Images were acquired using a Retiga R6 (Q-Imaging) camera. 6 randomly chosen positions per well were captured every 10 min. Individual cells were tracked using Manual tracking software (ImageJ) and Chemotaxis tool set (ImageJ).

### Inverted invasion assay

Inverted invasion assays were performed based on the protocol as described previously (Hennigan et al., 1994). Collagen I (rat-tail; final concentration 1.5 mg/mL; Gibco) supplemented with 25 μg/mL fibronectin was allowed to polymerize in inserts (Transwell; Corning) for 5 min at room temperature then a further 30 min at 37 °C. Transwells were inverted for wildtype MDA-MB-231 cells to be seeded directly onto the underside at a concentration of 5 x 10^5^ cells/mL and left for 4 h at 37 °C. Transwells were then re-inverted and placed in serum-free medium in the presence or absence of I3, afatinib (AFT), DANA or DMSO vehicle control, as indicated. Medium supplemented with 10% FCS and 100 ng/mL EGF, and the aforementioned inhibitors (or DMSO as control) was placed on top of the matrix, thereby providing a chemotactic gradient for invasion. After 72 h at 37 °C, all cells were stained 1 h at 37 °C with calcein-AM prior to imaging by confocal microscopy with an inverted Leica SP8 microscope using a 20× objective. Cells were considered invasive beyond 45 µm (red line marked in Fig 5H), and optical slices were imaged at 15 µm intervals. Invasion was quantified using the area calculator plugin in ImageJ, where the invasive proportion was obtained by measuring the fluorescence intensity of cells invading >45 μm divided by the total fluorescence intensity in all Z-stack images.

### Electron microscopy

MDA-MB-231 cells were grown to 40% confluency in 40 mm dishes. The next day, cells were incubated for 1 h, at 37 °C in serum-free medium, then for 6 min at 37 °C with serum-free medium containing 10 µg/mL of Gal3-HRP, anti-β1 integrin antibody (K20)-HRP or anti-⍰3 integrin antibody (ASC-1)-HRP in the presence or absence of 100 ng/mL EGF. Cells were put on ice and washed once with ice cold DMEM, 15 mM HEPES pH 7.3, 1% BSA, then twice with ice cold DMEM, 15 mM HEPES pH 7.3. Cells were incubated protected from light for 10 min on ice in freshly prepared 0.7□mg/mL 3,3’-diaminobenzidine (DAB) solution (Cat. no. D4293, Sigma) in DMEM, 15 mM HEPES pH 7.3, 1% BSA, followed by incubation for 20 min on ice with DAB, 30% H_2_O_2_ in the presence or absence of 10 mg/mL ascorbic acid (Cat. no. A0278, Sigma), always protected from light. Cells were washed three times for 10□min each on ice with DMEM, 15 mM HEPES pH 7.3, then fixed overnight at 4 °C in 2.5% glutaraldehyde (Cat. no. 16220, Electron Microscopy Sciences, Pennsylvania, USA) in 0.1□M Na-cacodylate (pH 7.2, Cat. no. 11653). The following day, cells were washed three times with 0.1 M Na-cacodylate for 10 min each, then post-fixed for 1 h at room temperature with 1% osmium tetroxide (OsO_4_) (Cat. no. 19152, Electron Microscopy Sciences) in 0.1□M Na-cacodylate, followed by one wash with 0.1□M Na-cacodylate for 10 min, and two washes with diH_2_O for 10 min each. Cells were then incubated with 4% UA for 45 min at room temperature, and rinsed three times with diH_2_O for 10 min each.

Tissue dehydration was performed at room temperature according to the following incubation protocol: 50% ethanol 5□min, 70% ethanol 5□min, 90% ethanol two times for 10□min each, 100% ethanol three times for 10□min each, followed by infiltration of Epon resin LX112 (Cat no. 21310, Ladd Research Industries, Vermont USA) at room temperature, 50% resin (LX112/EtOH 100% 1v/1v) for 30□min, then 100% resin LX112 for 1 h. Cells were embedded in capsules containing LX112 for 2-3 days at 60 °C. Samples were sectioned using an ultramicrotome (Reichert Leica UCT), and sections of 65□nm thickness were deposited on formvar/carbon-coated grids (form/carbon square 100 mesh – Cu: FCF100-CU-SB Electron Microscopy Sciences). Sections were imaged using a Tecnai Spirit electron microscope (FEI, Eindhoven, The Netherlands) equipped with a 4K CCD camera (EMSIS GmbH, Münster, Germany).

### Quantification of electron micrographs

A single slice was chosen for each cell, where the nucleus was clearly visible, roughly one quarter from the bottom of the cell. HRP-positive structures were counted and characterized as following: i) Clathrin or caveolin-coated vesicles (CCV), 80-120 nm in diameter with round shapes; ii) clathrin-independent carriers (CLIC), thin tubular structures of around 20 nm in diameter and up to 250 µm in length; iii) large vesicles or macropinosomes (MP), round structures with diameters above 120 nm. All structures were counted in each slice.

### Lattice light sheet microscopy (LLSM)

Genome-edited SUM cells were plated 2-4 h before imaging on 5 mm diameter #1.5 coverslips (Cat. no. 72195-05, Electron Microscopy Sciences). Cells were incubated for 2 min at 27 °C with 2 µg/mL anti-β1 integrin-Cy3 antibody or with 1 µg/mL transferrin-Cy3 in the presence or absence of 100 ng/mL EGF in LLSM imaging media (phenol red-free DMEM, high glucose, glutamax, supplemented with 1% BSA, 0.01% penicillin–streptomycin, 1 mM pyruvate, and 20 mM HEPES pH 7.3). The coverslips were transferred to the LLSM coverslip holder and inserted into the imaging chamber containing LLSM imaging media (for increased stability of the optical system). Imaging started within 5-8 min at 27 °C. LLSM was performed on a commercialized instrument of 3i (Denver, USA), as previously described (Chen et al., 2014). Cells were scanned incrementally through a 20 μm long light sheet in 600 nm steps using a fast piezoelectric flexure stage equivalent to ∼325 nm with respect to the detection objective and were imaged using two sCMOS cameras (Orca-Flash 4.0; Hamamatsu, Bridgewater, NJ). Excitation was achieved with 488 nm (Sapphire Coherent) or 560 nm diode lasers (MPB Communications) at 20% acousto-optic tunable filter transmittance with 100 mW (initial box power) through an excitation objective (Special Optics 28.6× 0.7 NA 3.74 mm water-dipping lens) and detected via a Nikon CFI Apo LWD 25× 1.1 NA water-dipping objective with a 2.5× tube lens with a final pixel size of 104 nm. LLSM imaging was performed using an excitation pattern of outer NA equal to 0.55 and inner NA equal to 0.493. Composite volumetric datasets were obtained using 10 ms exposure/slice/channel at a time resolution of 2.5 s per total cell volume (60 slices). Fifty time points were acquired within 2 to 3 min. Acquired data sets were analyzed using an adapted version of cmeAnalysis3D software published previously (Aguet et al., 2016; Renard et al., 2020). The software package can be found as part of the Github repository of LLSM tools in https://github.com/francois-a/llsmtools/. cmeAnalysis3D was implemented in Matlab 2021b. Quantification of LLSM images was performed on raw images (see Quantitative analysis of lattice light-sheet images). Napari, a□multi-dimensional image viewer for Python was used for 3D rendering (doi: 10.5281/zenodo.3555620).

### Quantitative analysis of Lattice light-sheet images

Post-processing of raw data volumes was carried out as described previously (Renard et al., 2020). Automated detection of AP2 structures or punctate structures of fluorescently-labeled cargoes in 3D was performed by numerical fitting with a model of the microscope point spread function (PSF) as described previously (Aguet et al., 2016). Automated tracking of cargo and clathrin-adapter protein AP2 was calculated using the u-track software package (Jaqaman et al., 2008), included in the cmeAnalysis3D software (Aguet et al., 2016). We used AP2 and cargo detection positions to map the membrane shape using the Matlab function of alphaShape (Renard et al., 2020). Subsequently, we estimated the displacement of each cargo from the membrane position orthogonally inwards of the cell. The distinction between punctate structures diffusing on the membrane and internalized ones, whether by clathrin dependent (AP2-positive) or clathrin independent (AP2-negative) was made by using the membrane position detection. An event was thereby counted as an endocytic event, if an object underwent a net displacement of at least 150 nm from the initial position inside the membrane proximal zone (Renard et al., 2020). The membrane proximal zone was limited by the contour of the alphaShape and a line 400 nm inwards of the cell. To calculate the number of AP2-positive and AP2-negative events, for every qualified internalization event of the cargo channel, the presence of AP2 was scored using the cmeAnalysis3D software (Aguet et al., 2016). Only tracks with a duration longer than 8 sec were used for the analysis. Calculation of lifetime distributions and intensity cohorts were performed as described in detail previously (Aguet et al., 2016). Quantification of LLSM images was performed on raw images. The raw LLSM images, used in Figures 5C-E and in Supplemental Movies 1-5 were denoised using ND-SAFIR (Boulanger et al., 2010), integrated into the BioImageIT platform (Prigent et al., 2022) before rendering in napari (doi: 10.5281/zenodo.3555620). Statistical analysis of Figure 5F was performed using Prism v9.4.1 software (Graphpad Inc).

### Animal procedures

All experiments were approved by the National Cancer Institute (National Institutes of Health, Bethesda, MD, USA) Animal Care and Use Committee and were compliant with all relevant ethical regulations regarding animal research. Twelve wildtype C57BL/6NCr mice were obtained from Charles Rivers Laboratory (Frederick, MD, USA) and divided in two cohorts. For each cohort, 3 mice were treated with afatinib and 3 with DMSO (vehicle). Briefly, stocks of 50 mg/mL afatinib (Medchem express, Princeton, NJ, USA) were dissolved in DMSO and stored at −80 °C. Afatinib or an equivalent volume of DMSO (control) was delivered as a single dose by oral gavage in 0.5% (v/v) methylcellulose and 0.2% (v/v) Tween-80 dissolved in deionized water. Six hours after the treatments, animals were euthanized by CO_2_-inhalation followed by cervical dislocation. The tongues were excised, and 1-2 mm of the anterior portions were cut and flash frozen in vials containing Ck Mix-lysis beads (Bertin Corp, Rockville, MD, USA). The remaining posterior parts of the tongues were covered with cryo-embedding media OCT (Sakura Finetek, Torrance, CA, USA), snap frozen on powdered dry ice, and stored at −80 °C until sectioning.

#### Gal3 and EGFR staining

Within 48 h of embedding, 5 μm coronal tongue sections were cut on a Leica CM1860 Cryostat (Leica Biosystems, Buffalo Grove, IL, USA), mounted onto charged superfrost-plus glass slides (Electron Microscopy Sciences, Hartfield, PA, USA) and immediately processed for labeling with Gal3. Sections were thawed at room temperature for 5 min, and OCT was removed by a brief 2 min wash in PBS. Sections were fixed for 10 min in 4% PFA in PBS followed by 3x 5 min washes in PBS. Sections were incubated for 1 h at room temperature in 5% BSA followed by staining overnight at 4 °C with CF594-conjugated recombinant Gal3 (200 nM = 5 µg/mL) or 1:100 anti-EGFR (#D38B1, Abcam) in PBS containing 1% BSA. The following day, sections were washed 2 times 5 min in PBS, incubated with 5 µg/mL DAPI for 10 min followed by two brief washes in PBS. Sections were mounted in Fluoromount G (Invitrogen, Carlsbad, CA, USA) and imaged using a Leica SP8 confocal microscope (Leica Biosystems, Buffalo Grove, IL, USA) equipped with a 40x objective captured at a 512×512 pixel size with the following white light laser and filter settings: DAPI: λex = 405 nm, λem = 416-441 nm; Alexa Fluor 568: λex = 575 nm, λem = 585-663 nm; Alexa Fluor 594: λex = 594 nm, λem = 604-637 nm; Cy5: λex = 650 nm, λem = 660-795 nm.

#### Image analysis

Two random areas of the epithelial layer were imaged for each tongue. Stacks of 40 optical sections (0.35 µm optical thickness) were acquired and converted into maximal projections view using the ImageJ2 (v2.9.0/1.53t) sum-projection plug-in. The lectin fluorescence intensity of the basal layer was quantified by measuring the mean intensity in a region of interest. Intensities were normalized for the mean intensity of vehicle-treated mice within each cohort. Normalized data from the two cohorts were pooled. Wilcoxon signed-rank test used for statistical analysis was conducted using R (4.2.1, 2022-06-23, R Core Team) and R Studio (2022.07.0, R Studio Team).

#### Immunoblotting of mouse tongues

400 µL RIPA buffer supplemented with 2x PhosSTOP and 2x complete protease inhibitor (Roche) were added to the flash-frozen tongue biopsies and lysis was performed using a Precellys 24 tissue homogenizer with Ck mix beads (Bertin Corp, Rockville, MD, USA) (3 pulses, 6.500 x g for 30 s/pulse). Debris were removed by centrifugation at 4 °C (21.000 x g for 10 min), supernatant isolated, and protein concentration quantified by DC protein assay (Bio-Rad, Hercules, CA, USA). Samples were diluted to 1 µg/µL in 1x Laemmli sample buffer (Bio-Rad, Hercules, CA, USA). 40 µg of protein per sample were separated on a 4-20% Criterion TGX gel (Bio-rad, Hercules, CA, USA). Subsequently, proteins were blotted onto a 0.4 µm PVDF membrane (EMD Milipore, Billerica, MA, USA) in a wet transfer system (20 V, overnight at 4 °C). Blots were blocked for 1 h at room temperature in 5% BSA in TBS-T, followed by overnight incubation at 4 °C in primary antibodies: Anti-tubulin (DMIA, 1:10,000, Thermo Fischer Scientific, Pittsburg, PA, USA) or anti-pEGFR (D7A5, 1:1,000, Cell Signaling Technologies). The following day, blots were washed 3 times 5 min in TBS-T and incubated for 1 h at room temperature with secondary antibodies: HRP-conjugated anti-rabbit IgGs (1:2,000, Cell Signaling Technologies) or HRP-conjugated anti-mouse IgGs (1:2,000, Bio-Rad, Hercules, CA, USA). Blots were washed 3 times 5 min in TBS-T and developed with SuperSignal West Pico PLUS chemiluminescent substrate (Thermo Scientific, Pittsburg, PA, USA) according to manufacturer’s protocol. Developed blots were imaged on a chemidoc system (Bio-Rad, Hercules, CA, USA).

## Supplemental Information Titles

Table S1. Gal3 and SNA interacting proteins

Table S2. Proteins that are significantly decreased in SNA pulldowns after EGF stimulation

Table S3. Proteins significantly enriched in Gal3 pulldowns after EGF stimulation

Table S4. Retrograde proteome of HeLa cells

Table S5. EGF-sensitive Gal3 interacting proteins found in the retrograde proteome

Table S6. Table of siRNAs, primers, and CRISPR guides that were used in this study

Movie S1. LLSM recording of β1 integrin endocytosis in the absence of EGF (related to Figure 5C-H)

Movie S2. LLSM recording of β1 integrin endocytosis in the presence of EGF (related to Figure 5C-H)

Movie S3. LLSM recording of transferrin endocytosis in the absence of EGF (related to Figure 5C-H)

Movie S4. LLSM recording of transferrin endocytosis in the presence of EGF (related to Figure 5C-H)

Movie S5. 2D movie of anti-β1 integrin (K20) antibody endocytosis imaged by LLSM

---

**Figure S1.**
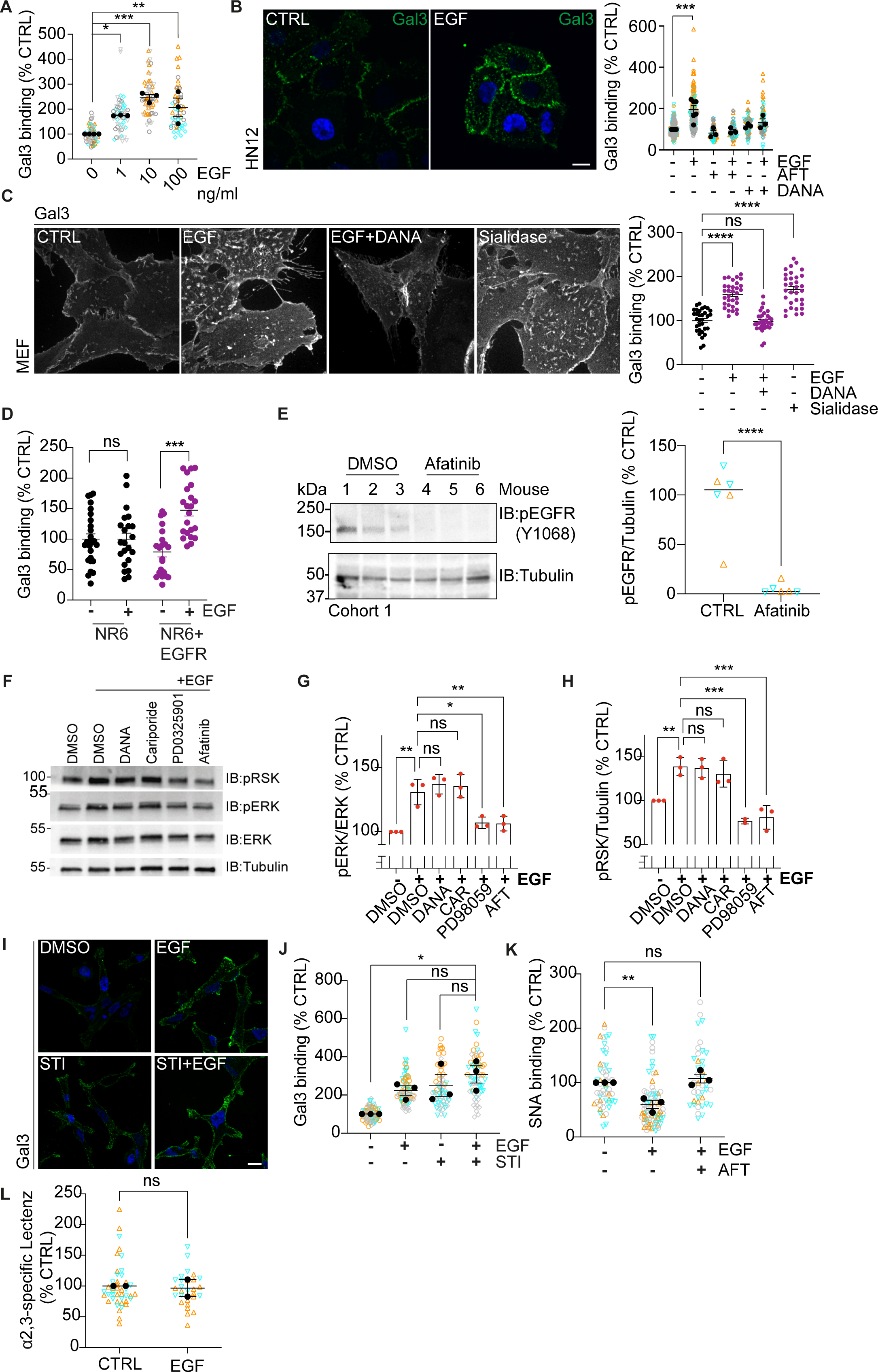
EGF effect on Gal3 binding to cells.

**Figure S2.**
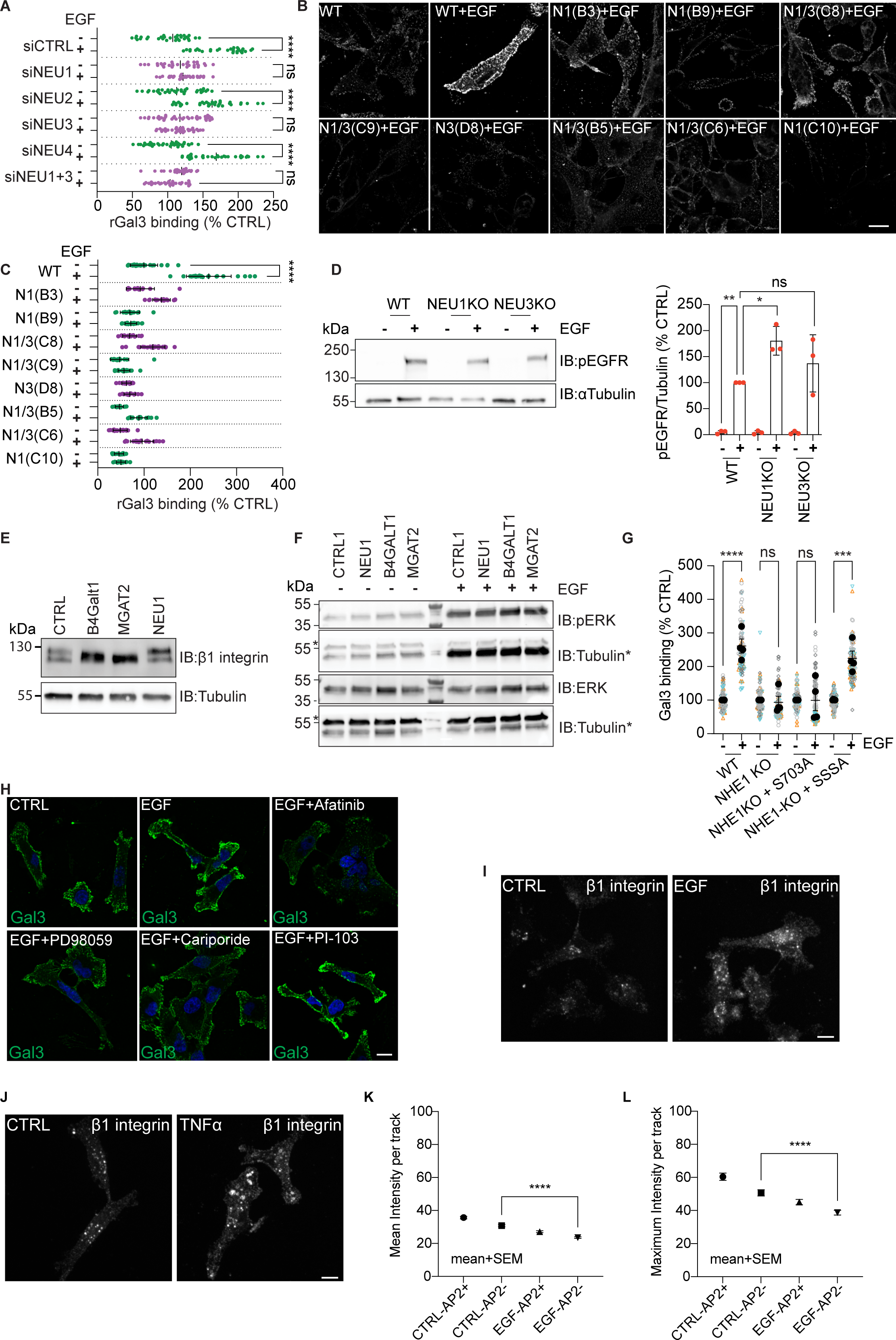
Neuraminidases and NHE1.

**Figure S3.**
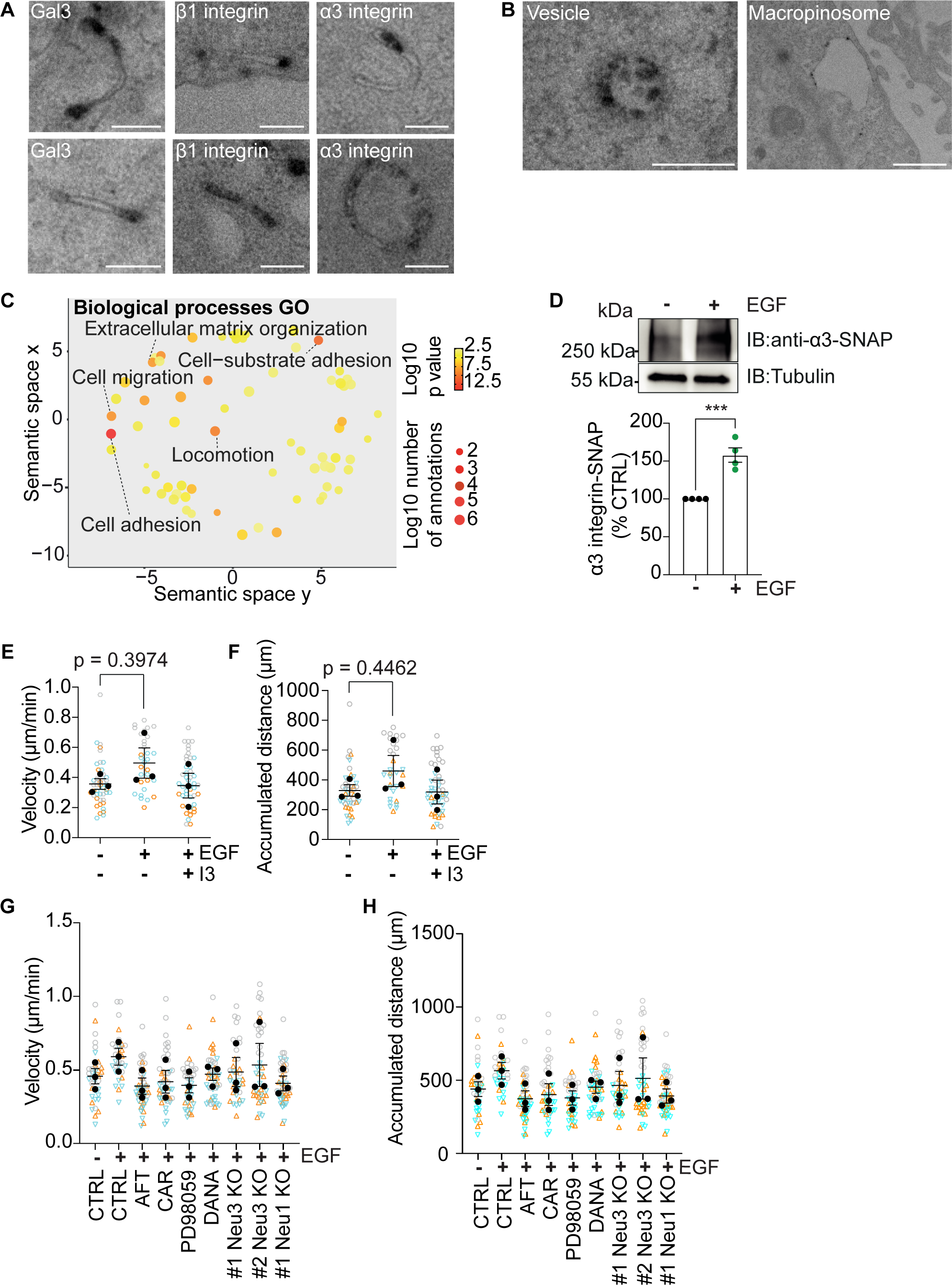
Cell migration.

