## Supplementary information for "Growth factor-induced desialylation for the fast control of endocytosis"

EGF-induced desialylation for the fast control of integrin endocytosis

**This pdf file includes:**

Figures S1 to S3

Tables S1 to S8

Movies S1 to S5

Figure S1. EGF effect on Gal3 binding to cells

(A) Serum starved MDA-MB-231 cells were incubated for 1 h on ice with Alexa488-labeled Gal3 in the presence of indicated concentration of EGF. Fluorescence intensity was determined for 4 independent experiments (≈ 100 cells per condition). One-way ANOVA with Dunnett’s multiple comparison test. (B) Gal3 (green) binding to HN12 cells. HN12 cells were incubated for 30 min at 37 °C with 200 nM AFT or 1 mM DANA in serum-free media before incubation for 1 h on ice in serum-free media with 200 nM Alexa488-Gal3 in the presence of absence of 100 ng/mL EGF. Total fluorescence/area from SUM confocal images was quantified from 6 independent experiments (≈ 200 cells per condition) and represented as %CTRL. One-way ANOVA with Dunnett’s multiple comparison test. Nuclei in blue. (C) Gal3 binding to mouse embryonic fibroblasts (MEFs). MEFs were serum starved for 30 min at 37 °C in the presence or absence of sialidase (1 unit) or 1 uM DANA. In the continued presence or absence of sialidase or DANA, the cells were then incubated for 30 min on ice with 200 nM Alexa488-Gal3 in the presence or absence of 100 ng/mL EGF. Total fluorescence/area from SUM confocal images was quantified from one representative experiment (≈ 30 cells per condition) out of 2, and represented as %CTRL. One-way ANOVA with Turkey’s multiple comparison test. (D) Gal3 binding to NR6 cells. NR6 cells (mutant EGFR) and NR6 expressing functional EGFR were serum starved for 30 min at 37 °C before incubation for 30 min on ice with 200 nM Alexa488-Gal3 in the presence or absence of 100 ng/mL EGF. Total fluorescence/area from SUM confocal images was quantified from one representative experiment (≈ 25 cells per condition) out of two, and represented as %CTRL. One-way ANOVA with Turkey’s multiple comparison test. (E) *In vivo* experiment in mice. Mice were treated for 6 h with 30 mg/kg AFT, tongues were excised and lysed in RIPA buffer. Lysates from all mice were probed with antibodies against pEGFR (Y1068) and tubulin. The quantification was normalized to first lane (cohort 1 orange triangle, cohort 2 blue triangles). (F) EGF-induced signaling. MDA-MB-231 cells were serum starved with the indicated inhibitors for 1 h at 37 °C before stimulation for 10 min at 37 °C in the continued presence or absence of the inhibitors and of 100 ng/mL EGF, as indicated. Cells were lysed in RIPA buffer, and 30 µg of lysate were probed with the indicated antibodies. (G,H) Quantification of 3 independent experiments as in (F) for the pERK signal (G) and the pRSK signal (H). One-way ANOVA with Tukey’s multiple comparison test. (I,J) General inhibition of sialylation. MDA-MB-231 cells were incubated for 72 h at 37 °C with 10 µM sialostatin (STI) or DMSO, serum starved for 30 min at 37 °C, and incubated for 1 h on ice with 200 nM Alexa488-Gal3 in the presence or absence of 100 ng/mL EGF. Total fluorescence/area from SUM confocal images was quantified from 3 independent experiments (≈ 75 cells per condition) and represented as %CTRL. One-way ANOVA with Holm-Šídák's multiple comparisons test. (K) EGF effect on SNA binding. MDA-MB-231 cells were serum starved for 1 h at 37 °C before incubation for 1 h on ice with 5 µg/ml of SNA-Cy5 in the presence or absence of 100 ng/mL EGF. Total fluorescence/area from SUM confocal images was quantified from 3 independent experiments (≈ 75 cells per condition) and represented as %CTRL. (L) EGF effect on Lectenz binding. MDA-MB-231 cells were serum starved for 1 h at 37 °C, incubated for 1 h on ice with 5 µg/ml of the α2,3-sialic acid linkage-specific Lectenz reagent in the presence or absence of 100 ng/mL EGF, fixed and labeled with neutravidin-488. Total fluorescence/area from SUM confocal images was quantified from one representative experiment (≈ 50 cells per condition) out of 2, and represented as %CTRL. Unpaired t-test.

For (E), the solid line indicates median of all data points. For all other quantifications, means ± SEM are shown. In (A,B,J-K), means from separate experiments are indicated by solid dots, and measurements of individual cells have different colored symbols for each experiment. ns = p > 0.05, ** p < 0.01, *** p < 0.001, **** p < 0.0001.

Figure S2. Neuraminidases and NHE1

(A) Neuraminidase screen. MEFs were transfected for 72 h at 37 °C with 100 nM siRNA against individual neuraminidases (25 nM of each oligo per sialidase). Post incubation, cells were serum starved for 30 min at 37°C, before incubation on ice with 5 µg/mL Alexa488-Gal3 in the presence or absence of 100 ng/mL EGF. Results of one experiment in duplicate. Data from 30 randomly picked cells in each set was analyzed and quantified. One-way ANOVA with Turkey’s multiple comparison test. (B) Screen of multiple clones for *NEU1*, *NEU3*, and *NEU1/3* CRISPR-edited MDA-MB-231 cells. Cells were serum starved for 1 h at 37 °C before incubation for 1 h on ice with 200 nM Alexa488-Gal3 in the presence or absence of 100 ng/mL EGF. (C) Quantification of total fluorescence/area from SUM confocal images from one representative experiment as in (B) are shown as %CTRL. One way ANOVA with Tukey’s multiple comparison test. (D) Autophosphorylation of EGFR. MDA-MB-231 cells were serum starved for 30 min at 37 °C before incubation for 10 min at 37 °C with 100 ng/mL EGF. Cells were lysed in RIPA buffer and Western blotting performed with anti-pEGFRY1068 antibodies. Quantification of 3 independent experiments as shown on the left. One-way ANOVA Tukey’s multiple comparison test. (E) Electrophoretic mobility of β1 integrin. Patient fibroblast harboring mutations in *B4GalT1*, *MGAT2* or *NEU1* were lysed, and lysates were analyzed by Western blotting with anti-β1 integrin antibodies. (F) pERK signaling. Patient fibroblast as in (E) were incubated or not for 10 min at 37 °C with 100 ng/mL EGF. Lysates of these cells were analyzed by Western blotting with the indicated antibodies. Note that pERK signals were increased by EGF treatment in all conditions. (G) Gal3 binding under NHE1 interference conditions. CRISPR-edited NHE1 knockout MDA-MB-231 cells and GFP-NHE1 mutant addback cells were serum starved for 30 min at 37 °C before incubation for 1 h on ice with 100 ng/mL EGF. Total fluorescence/area from SUM confocal images were quantified from 4 independent experiments and represented as %CTRL for each cell line. One-way ANOVA with Tukey’s multiple comparison test. (H) Incubation of serum starved MDA-MB-231 cells for 30 min on ice with 5.2 µg/mL Alexa488-Gal3 in the presence or absence of the indicated inhibitors and of 100 ng/mL EGF. (I,J) Effect of EGF and TNFα on the uptake of β1 integrin. MDA-MB-231 cells were serum starved for 30 min at 37 °C in the presence or absence of the indicated inhibitors and incubated for 10 min at 37 °C with 1 µg/mL of anti-β1 integrin antibody K20 in the continued presence or absence of the inhibitors and of EGF. Cells were then placed on ice and the cell surface bound antibody was removed by acid washes. After fixation, cells were labeled with Alexa488-coupled anti-mouse IgG antibodies. Quantification in Figure 4C. (K,L) Lattice light sheet microscopy imaging. Mean intensity (K) and maximum intensity (L) of β1 integrin endocytic tracks are shown from 3 independent experiments as in Figure 4G. 1802 tracks from 16 CTRL cells, and 2157 tracks from 18 EGF treated cells. Unpaired t-test. Note that both parameters decreased after EGF stimulation.

For all quantifications, means ± SEM are shown. ns p > 0.05, * p ≤ 0.05, ** p < 0.01, *** p < 0.001, **** p < 0.0001.

**Figure S3**. **Cell migration**

(A) Representative electron micrographs of CLICs containing the indicated HRP-coupled cargo proteins. Scale bars = 200 nm. (B) Representative electron micrographs of vesicular (left, scale bar = 200 nm) or macropinocytic structures (right, scale bar = 1 µm). (C) Cluster analysis of the summarized list of biological process gene ontology phrases associated with the list of 26 Gal3 binders that were significantly enriched after EGF stimulation. Spot size equals -LOG2 P value, spot color equals -LOG10 GO annotations. (D) Retrograde trafficking analysis (situation (i) in Figure 6A). HeLa cells stably expressing GalT-GFP-SNAP were continuously incubated for 4 h at 37 °C with benzylguanine-modified anti-α3 integrin antibody (ASC-1) in the presence or absence of 100 ng/mL EGF. The anti-α3 integrin antibody-SNAP conjugate in GFP-trap pulldowns on lysates from the indicated conditions of 4 independent experiments was quantified by Western blotting. Unpaired t-test. (E-H) Cell migration on cell-drived matrices. MDA-MB-231 cells were seeded on cell-derived matrices in 1% FCS and cell migration was monitored for 16 h at 37 °C in the presence or absence of indicated inhibitors. Velocity and accumulated distance were calculated from 3 independent experiments (≈ 50 cells analyzed per condition), using the manual tracking software ImageJ. One-way ANOVA with Dunnett’s multiple comparison test.

**Tables**

Table S1. Gal3 and SNA interacting proteins

MDA-MB-231 cells were subjected to pulldown with cell surface-bound SNA and Gal3, and identification by mass spectrometry. This table lists the 93 proteins common to both datasets.

Table S2. Proteins that are significantly decreased in SNA pulldowns after EGF stimulation

Mass spectrometry analysis of pulldowns with SNA that was bound to the surface of MDA-MB-231 cells in the presence or absence of EGF. List of 30 proteins whose binding to SNA was significantly decreased after EGF stimulation.

Table S3. Proteins significantly enriched in Gal3 pulldowns after EGF stimulation

Pulldowns with Gal3 that was bound to the surface of HeLa cells in the presence or absence of EGF. List denotes the 26 proteins whose binding to Gal3 was significantly enhanced after EGF incubation and proteins uniquely found in Gal3+ EGF condition.

Table S4. Retrograde proteome of HeLa cells

HeLa cells expressing GalT-GFP-SNAP were reacted with NHS-PEG9-BG versus DMSO control. List denotes all proteins that were significantly enriched in pulldowns from NHS-PEG9-BG modified cells.

Table S5. EGF-sensitive Gal3 interacting proteins found in the retrograde proteome

Comparison of proteins that were found enriched in binding to Gal3 after EGF stimulation and that were present in the retrograde proteome.

Table S6. Table of siRNAs, primers, and CRISPR guides that were used in this study

**Movies**

**Movie S1. LLSM recording of β1 integrin endocytosis in the absence of EGF (related to Figure 5C-H)**

Genome edited SUM159 cells expressing AP2-GFP (green) were incubated for 2 min with 2 µg/mL of anti-β1 integrin-Cy3 (magenta) antibody K20, and transferred to the LLSM for imaging at 27 °C. Full 3D volumes of 60 planes were acquired within 2.5 s. Colors indicate: In white, β1 integrin uptake tracks that were AP2-negative; in green, β1 integrin uptake tracks that were AP2-positive; in red, non-endocytic β1 integrin tracks.

**Movie S2. LLSM recording of β1** **integrin endocytosis in the presence of EGF (related to Figure 5C-H)**

Genome edited SUM159 expressing AP2-GFP (green) were incubated for 2 min with 2 µg/mL anti-β1 integrin-Cy3 (magenta) antibody K20 in the presence of 100 ng/mL EGF, and transferred to the LLSM for imaging at 27 °C. Full 3D volumes of 60 planes were acquired within 2.5 s. Colors indicate: In white, β1 integrin uptake tracks that were AP2-negative; in green, β1 integrin uptake tracks that were AP2-positive; in red, non-endocytic β1 integrin tracks.

**Movie S3. LLSM recording of transferrin endocytosis in the absence of EGF (related to Figure 5C-H)**

Genome edited SUM159 expressing AP2-GFP (green) were incubated for 2 min with 1 µg/mL transferrin (Tf)-Cy3 (magenta) and transferred to the LLSM for imaging at 27 °C. Full 3D volumes of 60 planes were acquired within 2.5 s. Colors indicate: In white, Tf-Cy3 uptake tracks that were AP2-negative; in green, Tf-Cy3 uptake tracks that were AP2-positive; in red, non-endocytic Tf-Cy3 tracks.

**Movie S4. LLSM recording of transferrin endocytosis in the presence of EGF (related to Figure 5C-H)**

Genome edited SUM159 expressing AP2-GFP (green) were incubated for 2 min with 1 µg/mL transferrin (Tf)-Cy3 (magenta) presence of 100 ng/mL EGF and transferred to the LLSM for imaging at 27 °C. Full 3D volumes of 60 planes were acquired within 2.5 s. Colors indicate: In white, Tf-Cy3 uptake tracks that were AP2-negative; in green, Tf-Cy3 uptake tracks that were AP2-positive; in red, non-endocytic Tf-Cy3 tracks.

**Movie S5. 2D movie of anti-β1 integrin (K20) antibody endocytosis imaged by LLSM**

Same conditions as Movie S1. This 2D movie shows a side view of 3 cell slices as maximum projection, perpendicular to the detection objective. Representative β1 integrin uptake events are shown: White arrows, AP2-negative; green arrows, AP2-positive. Scale bar: 5 µm.
